# The spatial landscape of progression and immunoediting in primary melanoma at single cell resolution

**DOI:** 10.1101/2021.05.23.445310

**Authors:** Ajit J. Nirmal, Zoltan Maliga, Tuulia Vallius, Brian Quattrochi, Alyce A. Chen, Connor A. Jacobson, Roxanne J. Pelletier, Clarence Yapp, Raquel Arias-Camison, Yu-An Chen, Christine G. Lian, George F. Murphy, Sandro Santagata, Peter K. Sorger

## Abstract

Cutaneous melanoma is a highly immunogenic malignancy, surgically curable at early stages, but life-threatening when metastatic. Here we integrate high-plex imaging, 3D high-resolution microscopy, and spatially-resolved micro-region transcriptomics to study immune evasion and immunoediting in primary melanoma. We find that recurrent cellular neighborhoods involving tumor, immune, and stromal cells change significantly along a progression axis involving precursor states, melanoma *in situ,* and invasive tumor. Hallmarks of immunosuppression are already detectable in precursor regions. When tumors become locally invasive, a consolidated and spatially restricted suppressive environment forms along the tumor-stromal boundary. This environment is established by cytokine gradients that promote expression of MHC-II and IDO1, and by PD1-PDL1 mediated cell contacts involving macrophages, dendritic cells, and T cells. A few millimeters away, cytotoxic T cells synapse with melanoma cells in fields of tumor regression. Thus, invasion and immunoediting can co-exist within a few millimeters of each other in a single specimen.

**STATEMENT OF SIGNIFICANCE:** The reorganization of the tumor ecosystem in primary melanoma is an excellent setting in which to study immunoediting and immune evasion. Guided by classical histopathology, spatial profiling of proteins and mRNA reveals recurrent morphological and molecular features of tumor evolution that involve localized paracrine cytokine signaling and direct cell-cell contact.

## INTRODUCTION

Tumorigenesis commonly involves a progressive failure of immune cells, particularly T cells, to detect cancer cells as they accumulate mutations promoting growth, invasion, and metastasis (1). The competition between editing by immune cells and escape by cancer cells generates a complex ecosystem whose molecular features and physical organization determine disease outcomes and responsiveness to therapy (2, 3). In the case of primary cutaneous melanoma, DNA sequencing has identified recurrent mutations in drivers such as BRAF, NRAS, PTEN, and TP53 (4–6) and dissociative single-cell RNA sequencing (scRNA-Seq) has revealed progression-associated changes in immune cell states (7). However, oncogenic transformation and immune escape remain only partly understood due in part to a high mutational burden in morphologically normal skin, estimated in Caucasians to be >100 driver mutations per cm^2^ by late middle age (8). Although treatment of metastatic melanoma has benefitted from modern targeted therapies guided by genetic biomarkers (BRAF and MEK inhibitors) and by immune checkpoint inhibitors, primary melanoma is treated surgically. It is diagnosed and staged using classical methods such as histopathological assessment of hematoxylin and eosin (H&E) stained formaldehyde-fixed paraffin-embedded (FFPE) skin biopsies, complemented in some cases by immunohistochemistry (IHC) (9).

Normal skin is characterized by evenly spaced melanocytes, which are neural crest-derived melanin-producing cells (10) located between cuboidal basal keratinocytes on the apical face of the dermal-epidermal junction (11). Fields of melanocytic atypia, the earliest signs of oncogenic transformation, involve increases in melanocyte number and density, enlargement, and irregularity of melanocyte nuclei, movement of melanocytes away from the dermal-epidermal junction (12), and loss of 5- hydroxymethylcytosine (5hmC) epigenetic marks (5, 13). These precursor fields can develop into melanoma *in situ* (MIS), a proliferation and confluence of malignant melanocytes within the epidermis but without invasion into the underlying dermis (14). MIS can spread within the epidermis and focally invade the superficial dermis without expansile growth, giving rise to radial growth phase melanoma, which has an excellent prognosis upon complete excision. However, invasive growth into the dermis is both expansile and highly mitotic, giving rise to vertical growth phase melanoma with a high potential for metastasis (15). Vertical growth phase melanomas can be endophytic or exophytic, corresponding to vertical growth down into the dermis or upwards above the skin, at times resulting in polypoid lesions that erupt from the surrounding skin (16).

The study of recurrent mutations found in cutaneous melanoma has yielded models of sequential tumor evolution starting with the formation of dysplastic nevi (4). However, while the removal of dysplastic nevi with higher grades of atypia is standard clinical practice (17) it is now thought that the majority of primary cutaneous melanomas are not derived from nevi, but rather arise de novo from fields of melanocytic atypia, particularly in sun-damaged skin (18, 19). The key features of these precursor fields, and the sequence of genetic events and immunosuppressive features that promote their progression to invasive melanoma remain poorly understood, as does the extent and impact of inter-patient and patient-to-patient variability. From a prognostic perspective, the depth of tumor invasion into the dermis (Breslow thickness) is a particularly important parameter (20) and is used in conjunction with the standard Tumor-Node-Metastasis (TNM) system used for melanoma staging (21). The number and locations of tumor-infiltrating lymphocytes (TILs) also have prognostic value (22). Finally, the Clark scoring system recognizes three distinct patterns for TILs: absent, non-brisk, and brisk (23). Absent describes both the absence of TILs and their failure to infiltrate tumor; non-brisk describes the restriction of TILs to scattered foci in the vicinity of the tumor, and brisk describes infiltration throughout vertical growth phase tumors or widely distributed along the invasive tumor front (24). In general, the greater the number of infiltrating TILs – the brisker the response – the more favorable the prognosis (25, 26). In some tumors, regions of inflammatory regression are also observed. In these regions T cells are observed to eradicate malignant melanocytes, leading to fields of fibrosis, vascular proliferation, and pigment incontinence, which are indicative of terminal regression (27). Inflammatory regression represents an example of successful and ongoing immunoediting but is currently incidental to diagnosis and of uncertain prognostic significance (28).

The great majority of studies on immune surveillance in primary and metastatic melanoma have involved either histologic analysis of H&E or IHC images, which are restricted to one to three markers per section, or sequencing of genomic mutations or mRNA profiling. However, several recent studies have demonstrated the potential for multiplexed imaging to provide greater insight into the spatially restricted tumor and immune programs in melanomas at different stages (29, 30).

Here, we focus on the molecular and morphological analysis of histologic features commonly found in primary melanoma. We focus on features used for diagnosis and treatment decisions in specimens containing multiple distinct stages of diseases. These include precursor fields, melanoma *in situ*, radial growth phase melanoma, and/or invasive vertical growth phase melanoma as well as regions of inflammatory regression. Specimens were acquired from the Brigham and Women’s Hospital dermatopathology tissue bank and, like virtually all primary melanomas, were available only in fixed form (FFPE) as a diagnostic necessity; only a few were subjected to or consented for DNA sequencing. The spatial organization of the tumor microenvironment (TME) was analyzed using 20 to 30-plex fluorescence microscopy (CyCIF) and either conventional wide-field microscopy or 3D optical sectioning followed by deconvolution (31). We also performed transcriptional profiling of selected micro-regions using two different methods for micro-region transcriptomics (mrSEQ: GeoMx and PickSeq) (32, 33). The resulting molecular and morphological data were then correlated with local histopathology as determined from H&E images by board-certified dermatopathologists. To preserve the spatial relationships of different histologies and to provide sufficient statistical power (34) CyCIF and H&E imaging were performed on whole slides, not tissue microarrays (TMAs) or small fields of view (FOVs) (35).

Using differential expression analysis and unsupervised clustering of mrSEQ data and spatial statistics on CyCIF data we identified molecular programs associated with histopathologic progression. In many cases, immunoediting by activated T cells was observed within a few millimeters of near-complete immune exclusion from invasive melanoma. Immunosuppressive niches were highly localized, in some cases only a few cells thick, and high-resolution imaging showed that they contained PDL1 expressing myeloid cells in direct contact with PD1 expressing T cells.

## RESULTS

### Multimodal profiling of spatially distinct regions within cutaneous melanoma

A total of 70 tissue regions (histological ROIs) with pre-cancer or cancer histologies were identified in eleven FFPE specimens of primary cutaneous melanoma, one locoregional metastasis, and one distant skin metastasis (specimens MEL1 to MEL13; **Supplementary Tables S1** and **S2;** histological features and annotations are described in **Supplementary Table S3**). Analysis of H&E-stained specimens by board-certified dermatopathologists confirmed the presence of one to five histological ROIs (average 2.4 per specimen) corresponding to precursor fields, melanoma in situ (MIS), invasive melanoma (IM), exophytic melanoma (EM), and inflammatory regression (IR) ∼5-20 mm apart from each other (summarized in **Supplementary Fig. S1A**). Serial FFPE sections (5 µm thick) were subjected to whole-slide, subcellular-resolution, 20-30 plex CyCIF imaging with different combinations of antibodies to generate complementary sets of image data (**Fig. 1A-1C, Supplementary Fig. S1A;** antibody panels described in **Tables S3, S4**). Antibodies included pan-cytokeratin (pan-CK) to stain keratinocytes in the epidermis; SOX10 and MITF to stain normal and atypical melanocytes and tumor cells (**Supplementary Fig. S1B**); smooth muscle actin (αSMA) to stain stromal cells, and CD31 to stain endothelial cells lining vessels. Immune cells were stained with lineage-specific cell surface proteins and functional markers (e.g., PD1) as described in **Supplementary Fig. S1C, Supplementary Tables S5,** and **S6**. Image analysis and data processing were performed using algorithms integrated into the open-source MCMICRO pipeline (36); staining intensities for lineage markers such as CD4, CD8, CD163, etc. were then binarized to distinguish among 13 immune cell types (**Fig. 1D-1F**).

**Figure 1:**
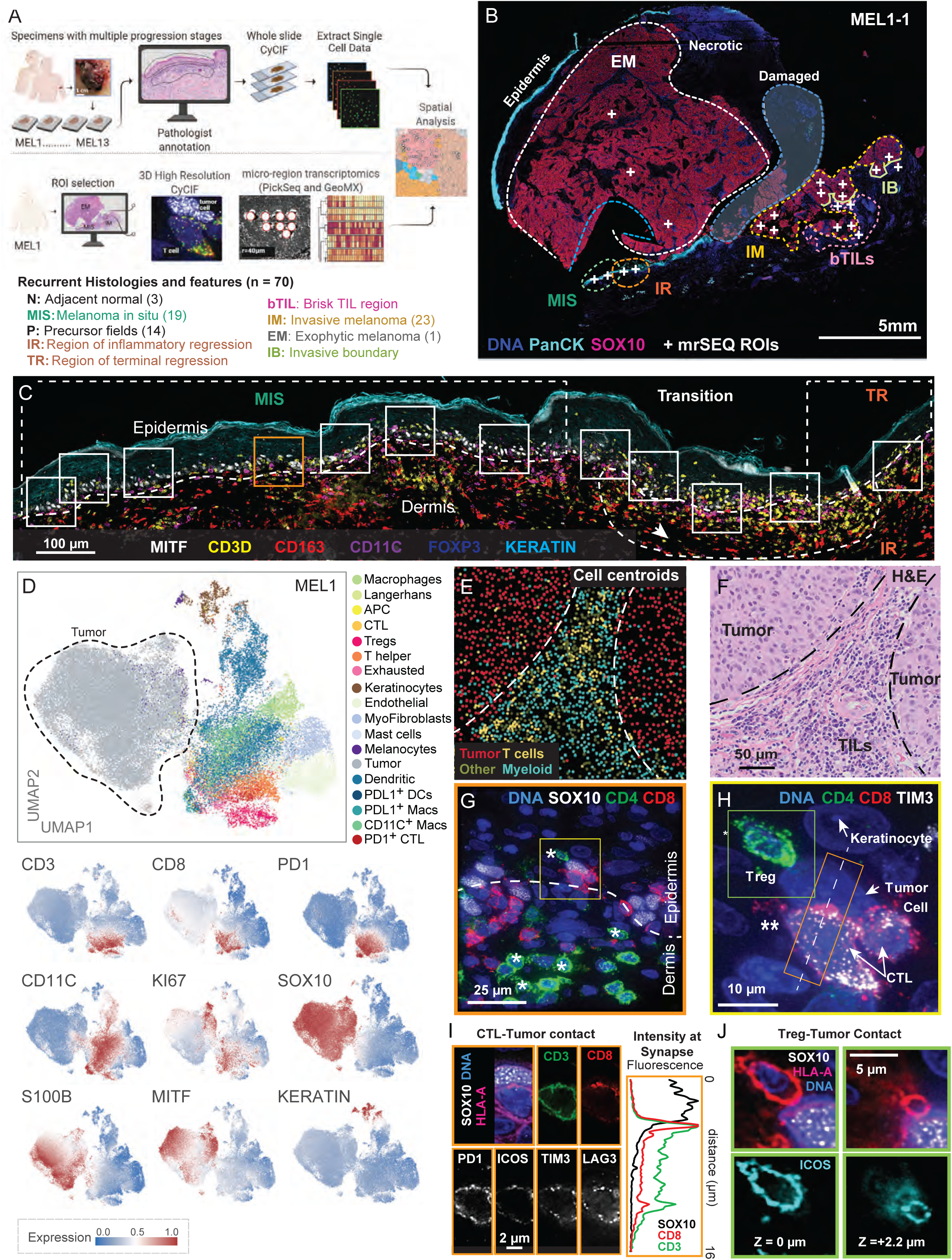
Multimodal profiling of cutaneous melanoma. **(A)** Conceptual framework of sample processing for cyclic immunofluorescence (CyCIF), high- resolution CyCIF, and micro-region transcriptomics: GeoMx and PickSeq (mrSEQ). Abbreviations for annotated histologies are shown below with color-coding used in subsequent figure panels. **(B)**) A 30-plex CyCIF image of a section of specimen MEL1-1 showing selected markers for epidermis (PanCK: cyan) and tumor cells (SOX10: red), highlighting annotated histologies and microregions (mROIs) that were subjected to mrSEQ (white +s). This specimen was likely torn during slide processing and thus, spatial arrangements in the region marked with a blue dashed boundary are not considered reliable. Other mrSEQ sites are shown in Supplementary Fig. 2A. **(C)** CyCIF image of MEL1-1 corresponding to the MIS and adjacent regions of inflammatory and terminal regression (IR and TR, respectively; outlined by dashed white lines). Rectangles depict the positions of 110 x 110 µm regions of interest (ROIs) in which high-resolution 3D deconvolution microscopy was performed. The region highlighted with orange is magnified in panel G. **(D)** Uniform manifold approximation and projection (UMAP) of single-cell data derived from CyCIF of patient MEL1 labeled by cell type (upper panel) and the signal intensities of individual markers (lower panels). Markers used for cell-type calls are shown in Supplementary Fig. 1C. The UMAP plot was built using 50,000 single cells that were randomly sampled from the full data set (n=1.1 x 10^6^). **(E)** Cell type assignments (with data points representing the centroids of cells) mapped to their physical locations in a portion of the bTIL region lying just beyond the IM in MEL1-1 **(F)** H&E image of the same region as in panel E. Regions of tumor and stroma are ”separated by dashed black lines. **(G)** A 21-plex high-resolution CyCIF image of a MEL1-1 MIS region (orange square in panel C) with selected markers shown as a maximum intensity projection staining for DNA (blue), tumor (SOX10: white), and T cells (CD4: green, CD8: red). The dermal-epidermal junction is denoted with a white dashed line and all FOXP3^+^ cells (as determined from other image channels; see Supplementary Fig. 1F) are denoted with an asterisk. Scale bar, 25 µm. Note that all images in panels G to J derive from a single multiplex CyCIF 3D image stack. **(H)** Magnified regions from panel G (outlined with a yellow box) showing staining of DNA (blue) and CD4 (green), CD8 (red), and TIM3 (white). Four cell types are labeled including a regulatory T cell (Treg, green box – shown in panel 1J) and two CD8^+^ CTLs interacting with a tumor cell (shown in panel I). The dashed line follows the axis of immune synapse polarization and gives rise to the intensity plot in panel I. The orange box depicts the locations of representative images in panel I. Scale bar, 10 µm. **(I)** Single optical section images of the immune synapse in panel H showing staining of tumor (SOX10: white), DNA (blue), and cell membrane (HLA-A: magenta) along with a series of single-channel images of functional T cell markers. The right panel shows the quantified spatial distribution of CD8 and CD3 along the dashed line in panel H. **(J)** Inset from panel H (outlined with a green square). Single optical section images of a tumor cell interacting with a Treg. Upper panels: staining for tumor (SOX10: white), cell membrane (HLA-A: magenta), and DNA (blue); lower panels: staining for Treg (ICOS: cyan). The two z-sections shown are spaced 2.2 µm apart.

More extensive molecular analysis was performed of specimen MEL1, which had the greatest number of distinct histologies (and spanned three tissue blocks MEL1-1, MEL1-2, and MEL1-3). MEL1 was an NF1-mutant, BRAF^wt^ tumor, which is one of four recurrent cutaneous subtypes identifiable in TCGA data (37). It was a large primary tumor involving both inward projecting vertical growth phase (nodular melanoma) as well as outward growing exophytic melanoma. A region of melanoma in situ (MIS) was co-extensive with regions of inflammatory and terminal regression in which immune editing had reduced or eliminated tumor cells; these regions contained dense infiltrates of CTLs, the majority of which were PD1^+^ (and thus activated) as well as Tregs. Invasive melanoma (IM) was located ∼10 mm away from the MIS and the invasive boundary (IB) of the nodular component had reached a depth of 4-5 mm and was surrounded by a domain of immune-rich stroma that was scored as a brisk TIL (bTIL) response. The patient from whom MEL1 was obtained developed loco-regional recurrence and distant metastases but was alive at the time of the last follow-up. MEL1 was characterized with a total of 80 different antibodies on five serial sections, subjected to micro-region transcript sequencing and 3D high-resolution imaging.

The ability of immune cells to make functional contacts with each other and with tumor cells is a fundamental feature of cancer immunoediting commonly quantified using spatial statistics (proximity analysis (38). In images collected at standard resolution (∼450 nm laterally), it is not possible to visualize the distinctive morphologies of immune synapses or PDL1 binding to PD1 (39). We, therefore, used 3D 21-plex CyCIF imaging with optical deconvolution on 110 µm square fields of view (FOV; ∼100 to 200 cells each) at a resolution of ∼220 nm laterally. Image stacks were collected from a total of 42 FOVs corresponding to regions of tumor invasion, MIS, and IR (where immune editing had reduced or eliminated tumor cells; **Fig. 1C** and **Supplementary Fig. S1D**). Among tumor and immune cells that were judged to be in proximity by proximity analysis of standard resolution images, we identified multiple examples of structures characteristic of functional cell-cell interactions.

Polarized interactions between the PD1 receptor and PDL1 ligand could also be imaged in this way (see below for data on ROIs and cell types). To estimate the frequency of such interactions, we performed a detailed inspection of two high-resolution FOVs lying at the tumor-stroma interface. A total of 199 cells (15 PDL1^+^ macrophages and 64 PD1^+^ T cells) were identified; in 58 cases these cells were judged to be within 20 µm of each other (a commonly used cutoff for proximity analysis) (40). In total, 21 immune cells (27%) had morphologies consistent with polarized PD1-PDL1 interaction. Thus, of the immune cells proximate enough to potentially interact directly, about one-quarter appeared to be involved in juxtracrine cell-cell interactions. These interactions were often complex, involving more than two cells. For example, **Fig. 1G** and **1H** show a SOX10^+^ tumor cell in contact with two CD8^+^ cytotoxic T lymphocytes and one CD4^+^ regulatory T cell (Treg; identified based on FOXP3^+^ staining in other imaging channels; **Supplementary Fig. S1E, F**), each of which was located at a different position on the tumor cell perimeter. Polarization of CD8 (a co-receptor for the T-cell receptor) at the site of contact between the tumor cell and one of the CTLs is consistent with the formation of an immune synapse. In this CTL, some TIM3 and LAG3 were partially localized to the synapse, although the majority of these proteins were sorted to the opposite side of the cell (**Fig. 1H, 1I** and **Supplementary Fig. S1H**). TIM3 and LAG3 are co-inhibitory receptors that function to regulate the activity of CTLs (41) and their presence on PD1^+^ CTLs showed that these cells are likely to be activated or possibly “exhausted”. The distribution of SOX10, CD3, and CD8 orthogonal to the plane of the cell-to-cell contact confirmed that the majority of CD8 (red line in the plot in **Fig. 1I**; **Supplementary Fig. S1G** and **S1I**) was found on the membrane of the CD3^+^ lymphocyte (green line) and approximately 500 nm away from the membrane of the adjacent SOX10^+^ tumor cell. Optical sectioning through the point of contact between the tumor cells and the Treg also revealed a contact (**Fig. 1J** and **Supplementary Fig. S1H**) that may be associated with the programming of tolerogenic activity.

Comprehensive characterization and quantification of cell-cell contacts detected by high-resolution tissue imaging await the development of better image recognition tools but our data provide clear and hitherto unavailable evidence that immune and tumor cells in close proximity to each other have structures characteristic of functional cell-to-cell contacts. Proximity analysis likely overestimates the frequency of these contacts whereas visual inspection of thin sections almost certainly results in an undercount because long processes perpendicular to the image plane are lost.

### Recurrent cellular neighborhoods associated with melanoma progression

To identify patterns of immune and tumor cell interaction that recur across patients and correlate with tumor progression, we used Latent Dirichlet Allocation (LDA) (42) (**Supplementary Fig. S2A**). LDA is a probabilistic modeling method that reduces complex assemblies of intermixed entities into distinct component communities (recurrent cellular neighborhoods; RCNs). LDA is widely used in biodiversity studies because it can detect both gradual and abrupt changes in the composition and arrangements of natural elements (cells in a tissue or trees in a forest) while effectively accounting for uncertainty and missing data (43, 44). To identify RCNs, ∼1.7 x 10^6^ single cells from MEL1-MEL13 were assigned to one of 12 basic classes based on the expression of cell type and state markers (e.g., proliferating, regulatory, exhausted) in 22-plex CyCIF data (**Fig. 2A** and **Supplementary Fig. S2B**). The data exhibited good signal to noise across critical markers and cell type assignment was robust to variation in gating (**Fig. 2B**). Across 70 histological ROIs annotated (regions of disease progression) we observed significant increases in percent of S100A+ SOX10+ cells between normal or precursor regions in comparision to MIS, and invasive melanoma, consistent with an increase in melanocyte-derived tumor cells (**Fig. 2C**) (45). S100 proteins are small calcium-binding proteins upregulated in melanoma and serum levels of S100B are used as a diagnostic marker of metastatic melanoma (although not a progression marker per-se; (46)). We trained spatial-LDA models using a 20 µm proximity radius so that RCNs would be enriched for cells in physical contact; latent weights were then clustered using k-means clustering (k=30) and grouped into ten informative meta-clusters (see methods). The generation of meta clusters made it possible to identify both direct and indirect interactions that recurred across the cohort. The RCNs corresponding to these meta-clusters were annotated based on cellular composition and frequency of occurrence in different ROIs and were then mapped to physical positions in the original specimens (**Fig. 2D**).

**Figure 2:**
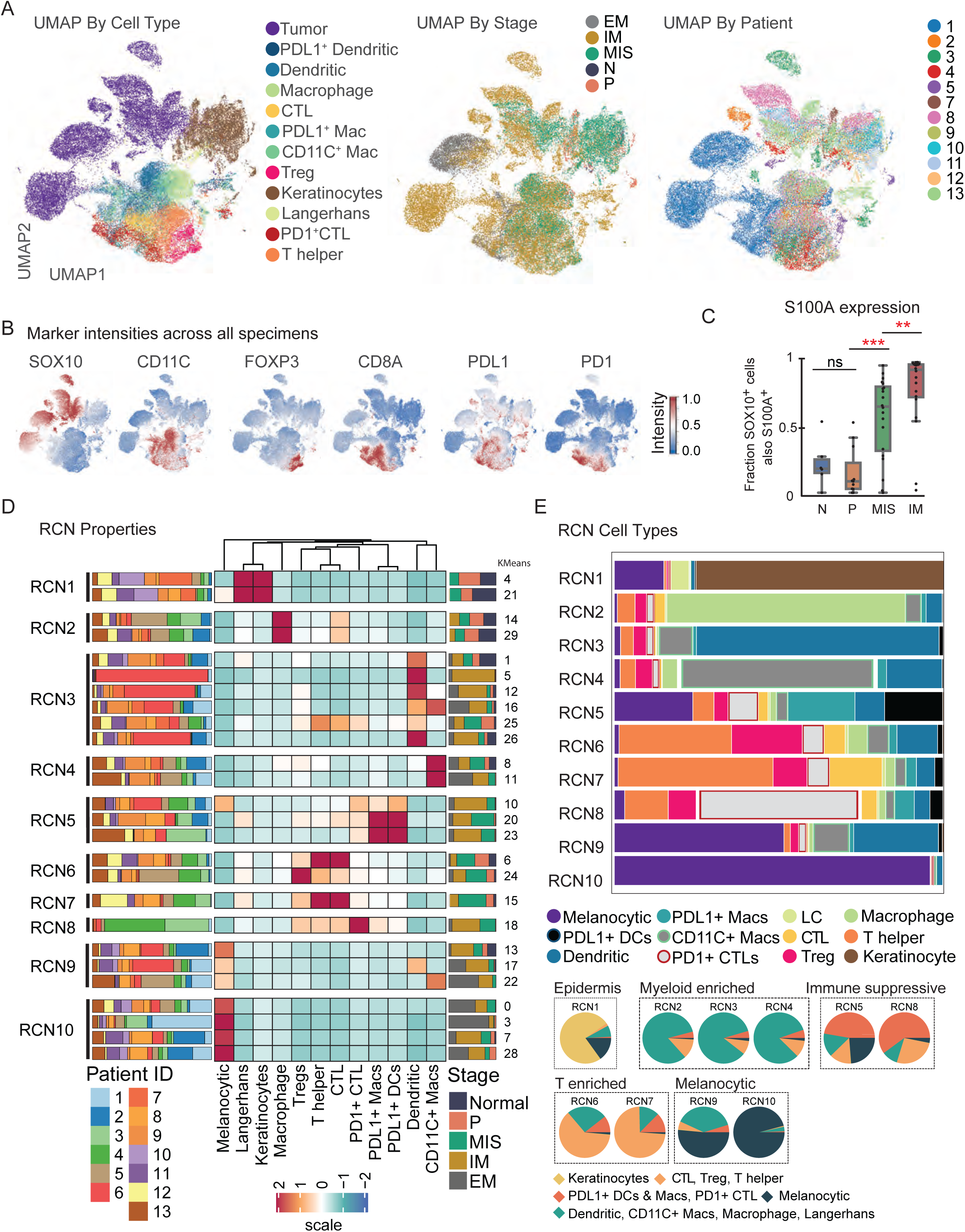
Recurrent cellular neighborhoods associated with melanoma progression **(A)** UMAP of single-cell data from 70 ROIs in 12 patients. The plot was generated using 50,000 single- cells that were randomly sampled from the full dataset of 1.5x10^6^ cells. The UMAP is colored based on the phenotype (left), disease progression stage (center), and patient ID (right). **(B)** UMAPs (shown also in panel A) representing feature plots of expression of selected protein markers. **(C)** The percentage of SOX10^+^ melanocytes or tumor cells expressing S100A within each stage of progression. **(D)** Heatmap showing the abundance of cell types within the 30 LDA-based cellular neighborhood clusters (numbers to the right of the plot); these were then reduced to the 10 meta-clusters (RCNs) shown to the left of the plot. The bar chart to the right of the heatmap depicts the distribution of progression stages within each cluster, and the bar chart to the left of the heatmap represents the distribution of patients within each cluster. **(E)** Bar plot depicting the detailed breakdown of cell-type proportions within each RCN (RCN1-10; x- axis). Pie charts depicting a simplified breakdown of cell types in each RCN; myeloid (green; dendritic cells, CD11C+ macrophages, macrophages, and Langerhans cells), lymphoid (light orange; cytotoxic T cell: CTL, regulatory T cells: Treg and helper T cell: T helper), immune-suppressive (dark orange; PDL1+ DCs, PDL1+ Macs, PD1+ CTL), melanocytes (dark blue) and keratinocytes (yellow).

Based on cellular composition, different RCNs corresponded primarily to epidermal, melanocytic, myeloid, T cell, and immune-suppressed populations (**Fig. 2E**). RCN1 was rich in keratinocytes (70% of the cells in this RCN) and Langerhans cells and was co-extensive with the epidermis (**Supplementary Fig. S2C**). RCN10 contained the largest number of cells (38% of all cells quantified), 90% of which were SOX10^+^; these corresponded primarily to tumor cells in regions of vertical growth phase melanoma (annotated as EM – exophytic melanoma, and IM – invasive melanoma) (**Fig. 2D**). In RCN10, tumor cells were densely packed together with few infiltrating cells (**Fig. 3A** and **3B**). In contrast, RCN9 (comprising ∼6.4% of all cells) contained equal numbers of SOX10^+^ and immune cells (36% and 34%, respectively) and corresponded to the interface between solid tumor and the dermis (red; **Fig. 2D, 3A-B**). Isolated pockets of RCN9 and RCN10 were also found in normal skin and in regions with adjacent melanocytic atypia and regions where SOX10^+^ cells clustered together (**Fig. 3C, Supplementary Fig. S3A**). The most abundant immune cells in RCN9 were CD11C^+^ macrophages and dendritic cells (80%) and the prevalence of this neighborhood increased significantly from precursor to MIS to invasive tumor, highlighting the formation of a myeloid-enriched tumor boundary (**Fig. 3D** and **Supplementary Fig. S3B**). When we quantified the proximity of tumor and CD11C^+^ myeloid cells using a 10 µm cutoff, we found that proximity volume scores increased from precursor to MIS to IM stages, independently confirming the observed increase in RCN9 frequency with progression (**Supplementary Fig. S3B, C**).

**Figure 3.**
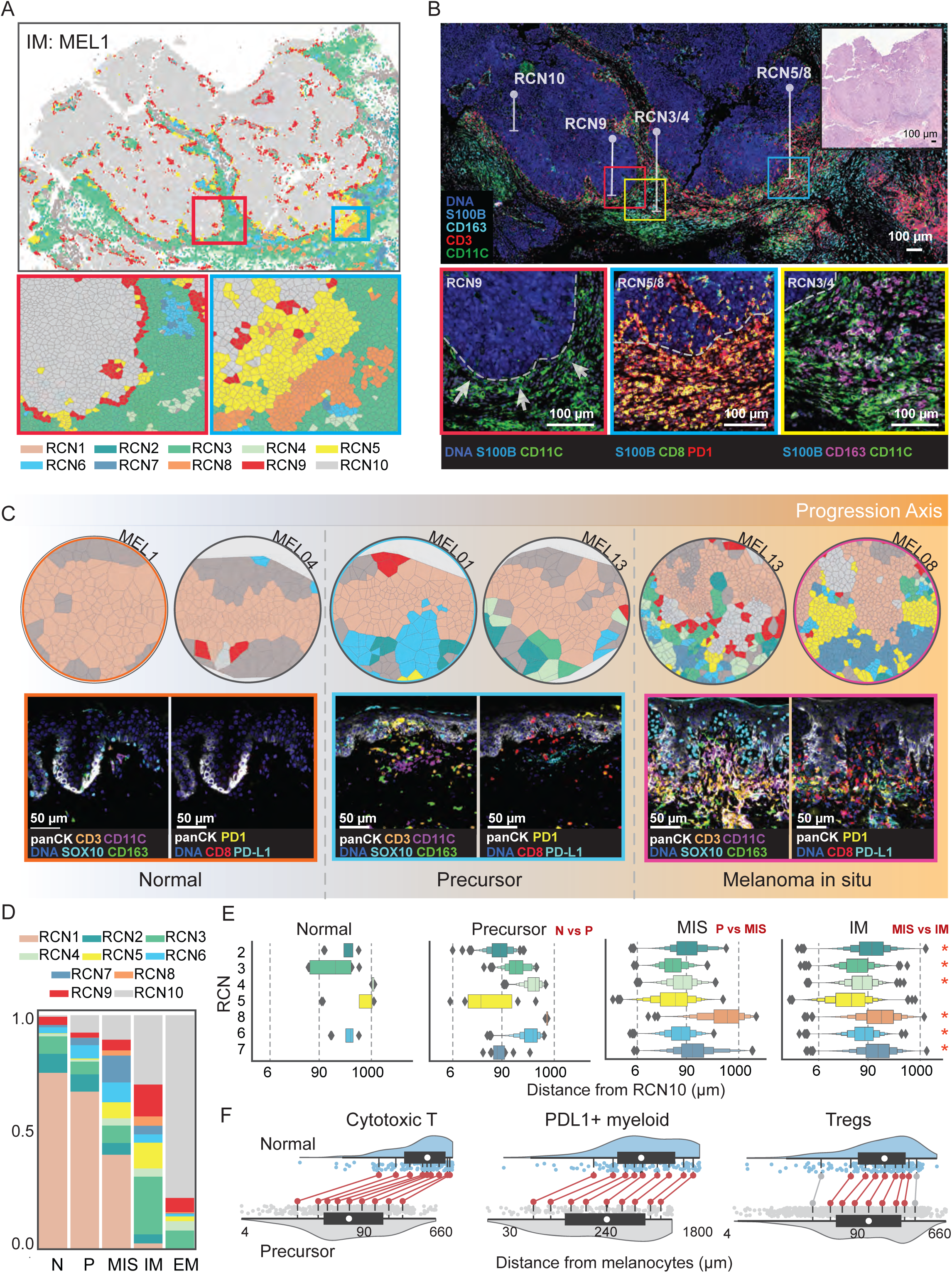
(A) Scatter plot (top) showing a field of view of the IM region (specimen MEL1-1). The cells are colored based on recurrent cellular neighborhoods (RCN1-10) that they belong to. The yellow and blue boxes represent regions that are magnified in the bottom panel (left and right, respectively) depicted as Voronoi diagrams. **(B)** Exemplary CyCIF images highlighting RCNs in the invasive front of specimen MEL1-1. The top panel shows an overall view of the invasive front stained for tumor cells (S100B: blue), macrophages (CD163: cyan), T cells (CD3: red), and dendritic cells (CD11C: green). The inset squares correspond to magnified panels at the bottom. H&E staining of a serial section of the same region is represented in the top right corner. The bottom left panel (yellow) highlights RCN9 enriched for dendritic cells (CD11C: green) at the tumor-stroma junction; the bottom center panel (blue) highlights RCN5/8 enriched with PD1^+^ CTLs (CD8: green; PD1: red) and bottom right panel (red) highlights RCN3/4 enriched with myeloid cells (CD163: magenta; CD11C: green). Scale bar, 100 µm; the dashed grey line represents the tumor-stroma boundary. **(C)** Voronoi diagrams of a representative field of views compiled from regions of N, P, and MIS. Each cell is colored based on the recurrent cellular neighborhood (RCN1-10) to which it belongs (as in panel A). Examples of corresponding CyCIF images from one patient in each case are provided at the bottom row. A magnified view is available in panel S3A. **(D)** Bar plot depicting the proportional distribution of RCNs (RCN1-10) among the disease progression stages (N, P, MIS, IM, and EM). **(E)** Box plots of the distribution of the shortest distance between cells in RCN 2-7 and RCN10 grouped based on progression stages. T-test (*P <0.05) depicts significant changes in mean distances between the compared stages. The comparison made is described on the upper right corner of each plot (e.g., N vs P). **(F)** Shift plot shows the distance between melanocytes and CTLs, PDL1^+^ myeloid cells, and Tregs in normal (top) and precursor (bottom) regions. Significance is calculated for each percentile (10, 20, 30, 40, 50, 60, 70, 80, 90) using the robust Harrell-Davis quantile estimator. Red indicates a significant difference (P <0.05) and grey represents non-significance for each percentile.

RCNs that were primarily made of immune cells could be subdivided into three classes: enriched for myeloid cells (RCN2-4), enriched for T cells (RCN6-7), and immune-suppressed (RCN5, 8). RCN2-4 contained overlapping sets of cells, with tissue-resident macrophages predominating in RCN2 and CD11C^+^ cells in RCN3 and 4 (**Fig. 2D**). RCN2 was found throughout the dermis (and had a distribution similar to that of tissue-resident macrophages) while RCN3 and 4 were found close to the invasive tumor (**Supplementary Fig. S3D**). RCN6 was rich in CD4^+^ T helper and Tregs and RCN7 was enriched for CTLs. RCN5 and 8 had high proportions of activated PD1^+^ CTLs as well as Tregs and PDL1^+^ myeloid cells, which are immunosuppressive (47). Five of the seven immune enriched niches (RCN3-7) significantly (P < 0.05) increased in frequency between precursor and MIS, while only one (RCN4) significantly increased between normal and precursor fields, reflecting recruitment of myeloid cells.

Two significant changes were observed between MIS and IM and this involved RCN9, which increased in abundance due to the formation of a PDL1^+^ sheath of cells at the IB, and RCN1, which fell in abundance due to the displacement of keratinocytes and proliferation of tumor cells in IM (**Supplementary Fig. S3E**).

When we quantified the proximity of immune rich RCNs (RCN2-8) to SOX10^+^ cells in RCN10 (i.e., melanocytes or tumor cells) we found that myeloid-enriched (RCN2, 4) and PDL1-enriched (RCN5) communities were significantly closer to RCN10 in precursor ROIs than adjacent uninvolved skin or later disease stages. In contrast, a cytotoxic community (RCN7) appeared closer to RCN10 in precursor samples than in MIS or IM (**Fig. 3E; Supplementary Table S7**). To confirm this finding, we measured the distance between melanocytic cells and the nearest PDL1^+^ myeloid cell or CTLs. We observed a significant decrease in distances for both cell types between normal and precursor stages. Tregs also showed a significant decrease in proximity to melanocytic cells in precursor fields (**Fig. 3F**). Thus, at the precursor stage, the recruitment of cytotoxic T cells was accompanied not only by immune resolution but also by the first signs of immunosuppression by myeloid cells. When RCNs were mapped back to the landscape of MEL1-1 (see **Fig. 1**), we found that the community of tumor cells near CD11C^+^ myeloid cells (RCN9) that were sporadically present in association with MIS had become a nearly continuous sheath at the invasive boundary of IM (**Fig. 3A-3C**). Immediately adjacent to the sheath of RCN9 cells we observed RCN3 and 4 myeloid niches in a mosaic pattern with RCN6 (T helper and Treg) and RCN5 (PDL1^+^ immunosuppressive) neighborhoods. The density of immunosuppressive niches was also highly variable even between nearby locations (**Fig. 3A** and **3B**). RCNs containing cytotoxic T cells (RCN7) and PD1^+^ CTLs (RCN8) were also intermingled, consistent with local activation of T cells. Moreover, whereas intermixing of tumor cells (RCN10) and multiple immune-rich RCNs was evident in MIS, in EM and IM the myeloid and immunosuppressive RCN (RCN5) was largely confined to areas immediately surrounding the CD31^+^ vasculature. Individual tumors differed in the specific arrangements of RCNs, and LDA generates statistical models subject to instance to instance variation, but it was consistent true that progression was associated not only with greater levels of invasion but also the formation of increasingly complex immune environments.

### PDL1-mediated immune suppression primarily involves myeloid not tumor cells

The importance of PD1-PDL1 interaction in melanoma is demonstrated by the success of anti-PD1 therapy. Across all 70 ROIs from 13 specimens, ∼70% of CTLs expressed the activation marker PD1 but we detected very few tumor cells expressing significant levels of PDL1, even in regions of tumor in which IFNɣ was highly expressed based on transcript profiling (see below; IFNɣ is a known inducer of PDL1). 3D deconvolution imaging proved to be more sensitive than conventional imaging in detecting PDL1, but even in MIS, in which immune and tumor cells were intermixed, only 5 of 106 tumor cells imaged at high-resolution in 12 FOVs were judged to be PDL1 positive. In these cases, imaging showed that PDL1 ligand on tumor cells and PD1 receptor on CTLs were co-localized, consistent with ligand-receptor binding (**Fig. 4A** and **Supplementary Fig. S4A**). In contrast to the paucity of PDL1^+^ tumor cells across all patients, significant co-occurrence (P < 0.05) was observed between PD1^+^ CTLs and PDL1^+^ macrophages and dendritic cells in 44 of 70 annotated histological domains; the frequency of this co-occurrence also increased with disease stage (**Fig. 4B**). To confirm that co-occurrence involved cell-to-cell interactions at least some of the time, we performed high-resolution 3D imaging of FOVs spanning the invasive front in MEL1-1 and observed frequent contact between PD1^+^ CTLs and either PDL1^+^ macrophages or dendritic cells with a concentration of PD1 and PDL1 at the site of cell-to-cell interaction (**Supplementary Fig. S4B** and **S4C**). In some cases, macrophages formed presumed inhibitory synapses with CTLs via cellular processes that extended at least one cell diameter (10 µm) from the macrophage (**Fig. 4C, Supplementary Video**), showing that non-adjacent cells can make functional contacts with each other. A substantial subset of PDL1^+^ myeloid cells also expressed TIM3, which is associated with immune suppression (**Fig. 4D** and **4E**).

**Figure 4.**
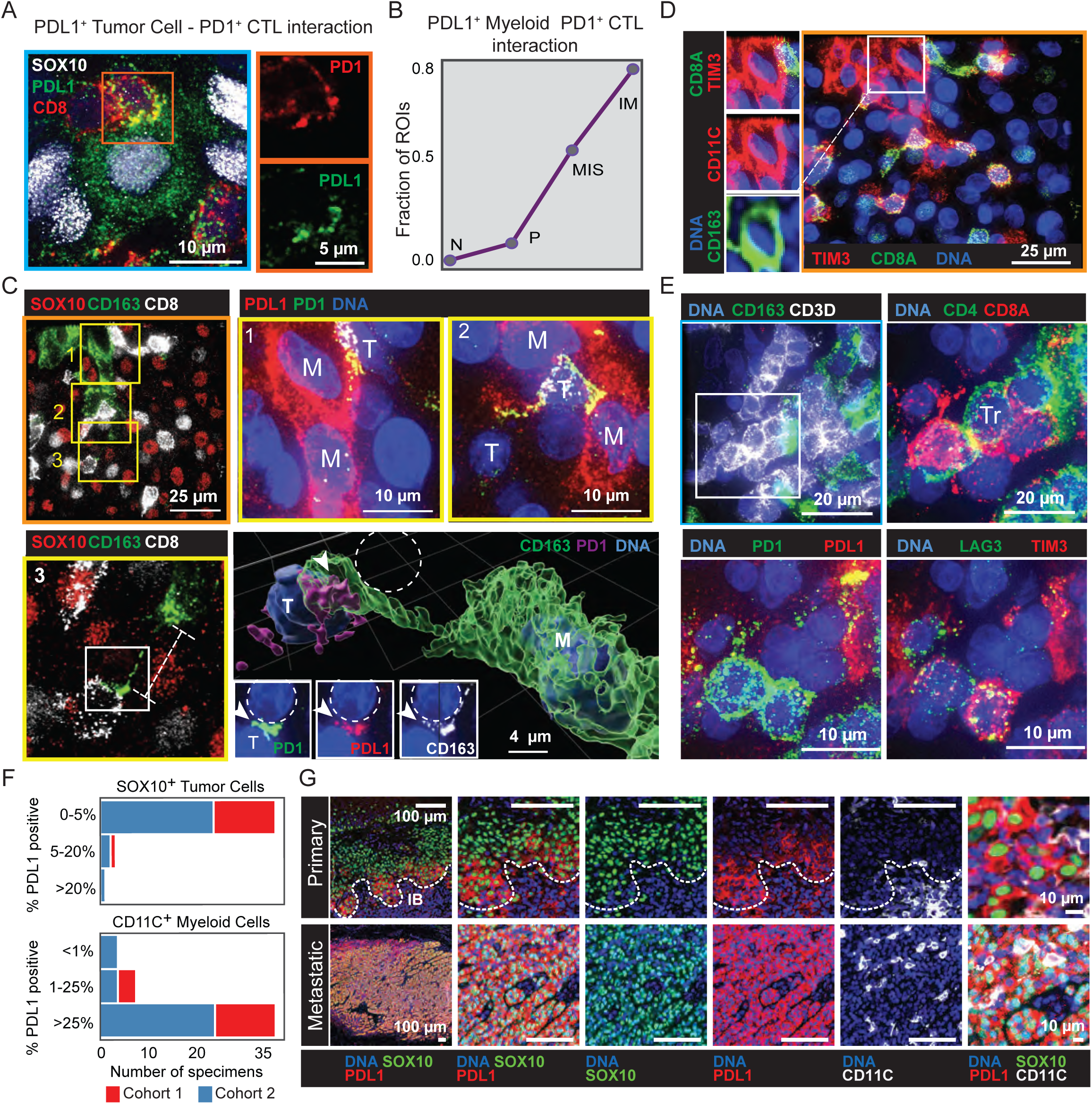
(A) Field of MIS from a whole slide CyCIF image of MEL1-1. A PDL1^+^ melanocyte (SOX10: white, PDL1: green) and CTLs (CD8: red) are being highlighted with an orange box (left panel). The right panel illustrates the polarization of PD1 (red) and PDL1 (green) to the point of contact between the interacting cells. Scale bar, 5 or 10 µm. **(B)** Line plot showing the percentage of ROIs that displayed significant (P <0.05) co-occurrence based on proximity analysis performed between PDL1^+^ CD11C+ CD163- dendritic cells and PD1^+^ CTLs. **(C)** Field of IM from a whole slide CyCIF image of MEL1-1 stained for tumor (SOX10: red), macrophages (CD163: green), and CTLs (CD8: white), with three fields of macrophage-CTL contacts (yellow boxes). Maximum-intensity projections imaged at high-resolution in fields 1 and 2 are stained for DNA (blue), PDL1 (red), and PD1 (green) with cells labeled as myeloid cells (M) and engaged T cells (T); field 3 shows tumor cells (SOX10: red), CTLs (CD8: white) and a macrophage (CD163; green). Inset white boxes in the bottom right panel show concentration of PD1 (red) and PDL1 (red) to the point of contact and the long connection between a macrophage (CD163: white) and a CTL is shown in a 3D reconstruction of the field 3. Scale bar, 25 µm, 10 µm or 4 µm. **(D)** Left panel shows the same CyCIF field of view as in panel C, stained for DNA (blue), TIM3 (red), and CD8 (green). The white inset box illustrates the staining of one CD163^+^ CD11C^+^ TIM3^+^ myeloid cell next to a CTL (right panel). Scale bar, 25 µm. **(E)** Maximum intensity projection from bTIL region (upper left panel) stained for DNA (blue), macrophages (CD163: green), and T cells (CD3D: white). The white inset is magnified and stained for T cell polarity (CD4: green, CD8: red), PD1-PDL1 axis (PD1: green, PDL1: red), and exhaustion markers (TIM3: red, LAG3: green). A Treg in this field is indicated with a label Tr. Scale bars, 20 and 10 µm. **(F)** PDL1 positivity in SOX10+ tumor cells (top) and CD11C+ myeloid cells (bottom). The proportions of PDL1+ tumor cells to all tumor cells (0-5%, 5-20% and >20%) and PDL1+ myeloid cells to all myeloid cells (<1%, 1-25% and >25%) are presented in both primary melanoma cohorts (cohort 1: MEL1-13 and cohort 2: 25 primary melanomas). **(G)** Fields of a primary melanoma and a melanoma metastasis from CyCIF images stained for DNA (blue), SOX10 (green), PDL1 (red), and CD11c (white). The upper panel shows an example of PDL1+ SOX10+ tumor cells at the deepest invasive region. PDL1+ metastasis is shown in the bottom panel. The tumor-stroma interface is indicated with a white dashed line. Scale bars, 100 and 10 µm.

We were surprised to find so few PDL1^+^ tumor cells in our specimens and therefore sought confirmation via analysis of an additional set of 25 primary melanomas. These specimens were annotated by dermatopathology as containing for radial (6/25) and vertical growth phase (16/25) histologies based on H&E images (as before) and subjected to low-plex immunofluorescence imaging for PD1, PDL1, SOX10 and CD11C followed by visual inspection of staining patterns by trained tissue biologists and pathologists. In these specimens PDL1^+^ SOX10^+^ tumor cells were abundant (estimated to be ≥20% positive) in only one specimen, present (5-20% positive) in two specimens, and infrequent (0-5% positive) in 22 of 25 specimens (**Fig. 4F** and **4G**; note that 5% threshold for PDL1 has previously been used to score tumor “PDL1 positivity” in melanoma) (48–50). In contrast >25% of CD11C^+^ myeloid cells scored as PDL1^+^ in 19 of 25 specimens. Moreover, using the same reagents, we have routinely found that nearly all metastatic melanomas contain abundant PDL1^+^ SOX10^+^tumor cells (51). When the three melanomas containing relatively abundant PDL1^+^ tumor cells were examined further, we found that PDL1-positivity was strongly enriched at the IB of vertical growth phase melanoma, which is enriched in cytokines secreted by immune cells (e.g. IFNγ) and the probable site of metastasis formation. We conclude that in the primary melanomas we imaged (n = 33 of 36 in total), the cells most expressing PDL1 were dendritic cells and/or macrophages, not tumor cells.

Recent data from the MC38 murine syngeneic model of colorectal cancer suggests that dendritic cells, not macrophages, may also be the relevant myeloid cell type for PDL1-mediated immunosuppression of activated CTLs in colonic adenocarcinomas (52). However, whereas the murine tumors analyzed by Oh et al. (52) contained many more PDL1^+^ macrophages than PDL1^+^ dendritic cells, we found that these two types of myeloid cells were similar in abundance in primary melanoma (1.2 to 1.4% of all cells). By high-resolution imaging of the invasive front, we also found multiple fields in which tumor cells, CTLs, dendritic cells, and other immune cell types (a subset of which expressed PD1 or PDL1) were all in direct contact with each other as part of extended networks (**Fig. 4E**). The presence of multi-dentate cell interactions and extended cellular processes containing immune-regulatory molecules suggests that multiple different immune cell types might communicate with each other via cell-cell contacts as well as autocrine or paracrine signaling. A more complete understanding of these interactions awaits high-resolution 3D reconstructions of the TME.

### Single-cell analysis of invasive tumor reveals large scale gradients in lineage, immune, and proliferation markers

Because LDA detects discrete differences between cells (most commonly in immune differentiation markers), it is insensitive to qualitative differences between cells of a single type. To quantify such differences - specifically in tumor cells - we used principal components analysis (PCA), and shift-lag analysis, focusing on cells in the invasive tumor (∼5 x 10^5^ malignant single cells) (**Supplementary Fig. S1A**). Principal components 1 and 2 (PC1 and PC2) explained 40% of the variance in these data, which represents good performance for a PCA model. The top loadings in score plots were KI67, the S100A and S100B proteins, and the MITF transcription factor (**Supplementary Fig. S5A**). KI67 is widely used to measure proliferation (55) and MITF is a master regulator of melanocyte differentiation (56). MITF is both a melanoma oncogene (57) and a determinant of drug resistance (58): an MITF^low^ state has been associated with de-differentiation and resistance to RAF/MEK therapy (59). Across the whole tumor, S100A, S100B, and MITF exhibited striking gradients in expression levels on short and long length scales (∼100-3000 µm), with the highest protein levels at the invasive margin, and lowest in the middle of the EM (**Fig. 5A, 5B** and **Supplementary Fig. S5B**). Thus, whereas clustering of sequencing data (7) emphasizes the presence of dichotomous MITF or S100 high and low states, imaging reveals continuous changes in protein levels through space. Spatial gradients involving morphogens have been widely studied in tissue development (60) but infrequently in cancer (61).

**Figure 5:**
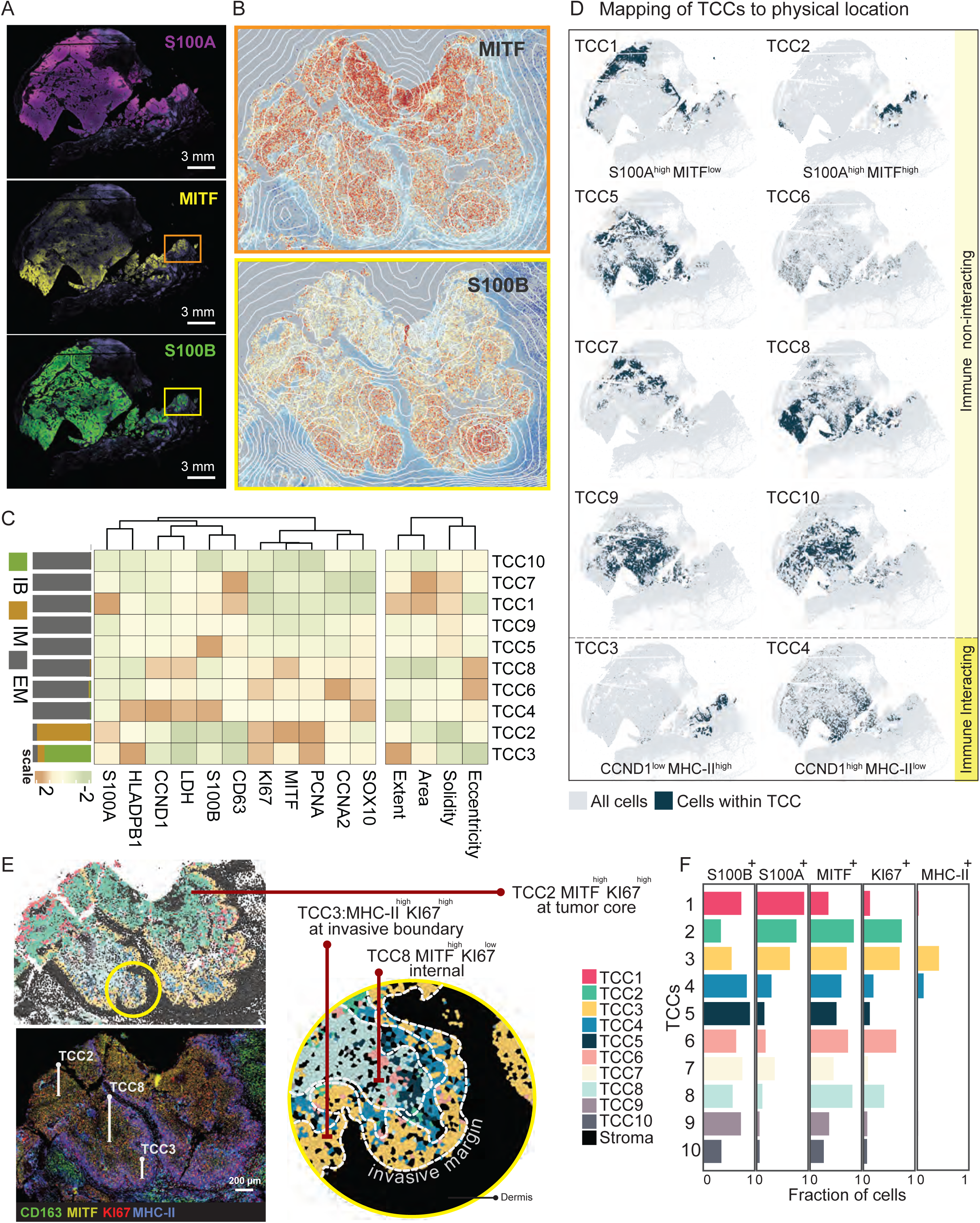
Single-cell analysis of invasive tumor. **(A)** CyCIF images of MEL1-1 stained for S100A (top panel), MITF (middle panel) and S100B (bottom panel). Boxes represent regions highlighted in panel B. Scale bars, 3 mm. **(B)** Insets from panel A of tumor region (IM) showing gradient expression patterns for MITF (top panel) and S100B (bottom panel). Contours describe averaged quantified marker expression. **(C)** Heatmap showing median expression of protein markers identified within TCC1-10 tumor cell communities. The bar plot on top of the heatmap shows the proportional estimate of the TCCs within histological annotations (EM, IM, or IB). The heatmap at the bottom shows the properties related to the shape of the cells (area, solidity, extent, and eccentricity) derived from the segmentation masks. **(D)** Scatter plot mapping the physical location of the derived tumor cell clusters (TCC1-10: dark blue) in MEL1-1. Each subplot represents the location of cells within a tumor cell community and other cells in grey. **(E)** Scatter plot (left panel) showing a field of view of the IM region. Cells are colored based on their tumor cell community (TCC1-10). The yellow circle highlights the region in panel B. The right panel is a CyCIF image of the same field of view (from specimen MEL1-1) stained for CD163 (green), MITF (yellow), KI67 (red), and MHC-II (HLADPB1: blue). Scale bar, 200 µm. Voronoi diagram (right panel) generated from a field of view at the apex of the invasive front (inset highlighted in yellow). Cells are colored based on the tumor cell community (TCC1-10) that they belong to. **(F)** Bar plots showing the percentage of S100B, S100A, MITF, KI67, and MHC-II (HLADPB1) positive cells within each tumor cell community (TCC1-10).

Spatial lag is a common spatial statistic in geography and ecology (62) and we used it to identify recurrent tumor cell communities based on continuous differences in protein levels. Clustering of spatial lag vectors revealed the presence of 10 tumor cell communities (TCCs; see methods for details of clustering; **Fig. 5C** and **5D**) that differed from each other in hyperdimensional features (combinations of markers) although in a few cases, single markers were dominant: MHC-II positivity for TCC3 and a MITF^high^ KI67^low^ state for TCC8 (**Fig. 5C** and **5D**). TCC1 corresponded to an S100A^high^ MITF^low^ pattern that was primarily found in EM, while MITF^high^ cells in TCC2 were primarily found in IM. TCC3 and TCC4 were either MHC-II ^high^ or CCND1^high^ and had distinctive spatial localization (**Fig. 5C** and **5D**).

The component of the IM facing the dermis had seven distinct TCCs each of which was 2-5 cell diameters thick. For example, TCC3 and TCC4 were found at the invasive boundary and significantly co-localized (P < 0.05 by co-occurrence analysis) with immune cells. TCC8 was found internal to TCC3 and TCC4, and TCC1 and TCC2 were primarily found internal to this, at the trailing edge (**Fig. 5E**).

H&E imaging has previously suggested that vertical growth phase melanoma might have the layered arrangement of tumor cells states revealed by spatial lag analysis of CyCIF data (63).

The invasive state of cutaneous melanoma cells is thought to involve an MITF^low^ slowly-cycling state (58). However, we found that the TCC2 community at the invasive front was comprised of 70 to 85% MITF^high^ KI67^high^ cells (**Supplementary Fig. S5D** and **S5E**). Further evidence of proliferation was provided by positive staining of many cells in this TCC with antibodies against cyclin A2, cyclin B1, phospho-Rb (pRB, which is highest in S-phase), and phospho-histone H3 (pHH3 a marker of mitosis; **Supplementary Fig. S6F**) with the highest rates of proliferation in IM (∼3-fold fewer proliferating cells were present in EM; **Supplementary Fig. S6E** and **S6F**). A second previously described feature of invasive melanoma is upregulation of genes involved in EMT in epithelial cells (in the case of melanoma these have been referred to as EMT-associated genes; (64, 65)) and anti-apoptotic programs (66); we observed both in tumor cells in IM (**Fig. 6G** and **Supplementary Fig. S6B**). Thus, the cells at the invasive boundary of MEL1 have several molecular properties previously associated with invasion, but they are neither MITF^low^ nor slowly proliferating (relative to the rest of the tumor). NGFR (CD271) and the AXL receptor tyrosine kinase are two other proteins widely studied for their roles in state switching and drug resistance in metastatic disease (67). However, we detected only sporadic NGFR expression in MEL1 tumor cells by either mrSEQ or imaging. AXL was detected only on the plasma membranes of keratinocytes and immune cells, not tumor cells (68). Thus, the primary melanomas we imaged differed in MITF, NGFR, and AXL expression from the melanoma cell lines used in most laboratory studies, most of which were derived from metastatic disease (69).

**Figure 6:**
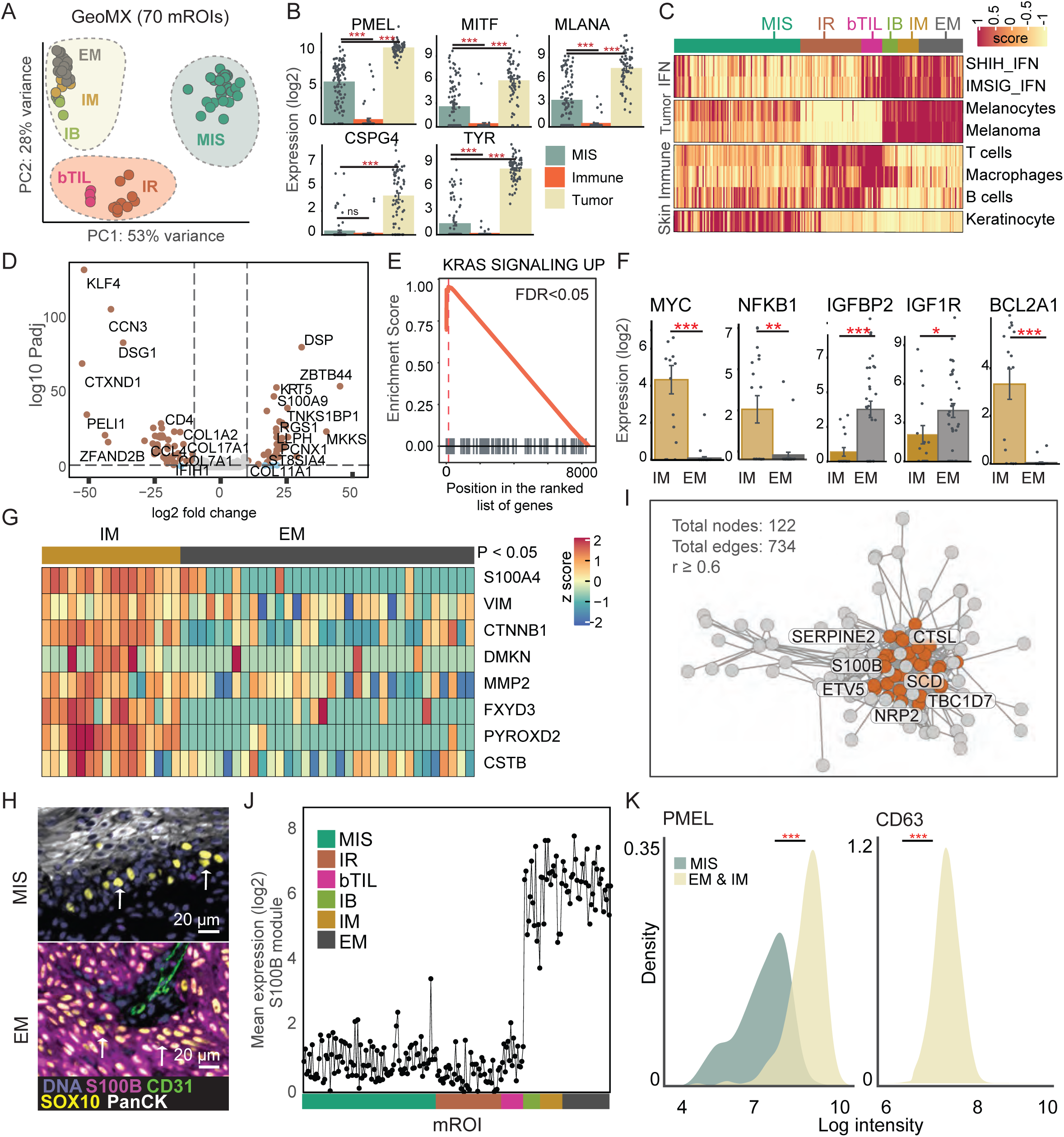
Micro-regional transcript profiling **(A)** Principal component analysis (PCA) plot of melanoma mrSEQ transcriptomes (GeoMX). Colors indicate regional histopathology: brisk TIL (bTIL: pink), inflammatory regression (IR: brown), MIS (green), invasive front (IB: light green), exophytic melanoma (EM: grey), and center of invasive melanoma tumor (IM: yellow). EM and IM are enriched for tumor cells in this analysis and IB contains mostly tumor cells with marginal immune infiltration. **(B)** Expression of selected melanoma-related marker genes in mrSEQ data (PickSeq) split into three broad groups based on the PCA of GeoMx data (panel A). Data is mean ± SEM. ***P <0.001; ns = not significant. **(C)** Single-sample gene set enrichment analysis (ssGSEA) on mrSEQ data (PickSeq). ssGSEA scores highlight enrichment of melanoma-related gene signatures in tumor mROIs (primarily IB, IM, and EM) and immune-related signatures in the immune-rich mROIs (IR, bTIL). **(D)** Fold-difference (log2) and significance (log 10 Padj) for expression of 19,500 genes between EM (n=34) and IM (n=16) mROIs (Pick-Seq). DEGs above (brown) and below (blue/grey) a significance threshold (P-adjusted = 0.05) and above a fold change threshold (log2 fold change = 10) are indicated. **(E)** GSEA for upregulation of KRAS pathway in IM (n=16) compared to EM (n=34) mROIs (PickSeq). FDR < 0.05. **(F)** Expression (log2) of MYC, NFKB1, IGFBP2, IGF1R, and BCL2A1 in IM and EM mROIs (PickSeq). Data is mean ± SEM; *P <0.05, **P <0.01, ***P <0.001. **(G)** Heatmap showing expression of genes (listed on the y-axis) known to play a role in epithelial to mesenchymal transition (PickSeq). All genes showed a significant difference between their mean expression in IM vs. EM mROIs (P <0.05). **(H)** CyCIF image showing a field of view in MIS (top panel) and EM (bottom panel) regions. The tissue is stained for melanocytes (SOX10: yellow), endothelial cells (CD31: green), keratinocytes (PanCK: white), and tumor cells (S100B: magenta). Arrows mark examples of melanocytes and tumor cells. Scale bar, 20 µm. **(I)** Correlation network sub-graph genes associated with S100B expression in mrSEQ data (PickSeq). Nodes represent genes, and the edges correspond to the correlation between them. Brown nodes represent the genes that belong to the S100B module. Selected genes are annotated. **(J)** Mean expression of 35 genes identified within the S100B module in mrSEQ data (PickSeq). The X- axis represents the mROIs grouped into the histopathological annotation category from which they were isolated. **(K)** Density plots illustrating the log scaled protein expression of PMEL and CD63 in MIS and tumor (EM&IM) regions imaged with CyCIF. ***P <0.001.

### Micro-regional transcript profiling identifies spatially distinct immune, mitogenic, and survival programs

To study the transcriptional programs associated with different immune neighborhoods and tumor cell communities that we identified using LDA and spatial lag analyses, we performed micro-region transcriptomics (mrSEQ) on a total of 292 microregions of interest (mROIs) of specimen MEL1 using PickSeq (32), which recovers 5-20 cells per 40 µm diameter micro-region of interest (ROI) and GeoMX (a commercial technology) which recovers ∼200-400 cells per ∼200 µm diameter mROI; **Fig. 1B** and **S6A**) (70). PCA of mrSEQ data revealed three primary clusters corresponding to (i) MIS, (ii) malignant tumor (EM plus IM), and (iii) regions of active immune response (IR – which were adjacent to the MIS and a bTIL region adjacent to the invasive boundary) (**Fig. 6A**). We found that markers commonly used to detect and subtype malignant melanoma (PMEL, MLANA, TYR, MITF, and CSPG4) were all strongly and consistently expressed at the gene level in mROIs from tumor domains (EM and IM), sporadically in MIS and not in immune-rich regions (IR, bTIL) confirming the annotation of these regions and the selectivity of the method (**Fig. 6B;** gene names are listed in **Supplementary Table S6**). Single-sample gene set enrichment analysis (ssGSEA) confirmed high enrichment of melanocyte signatures in tumor but not in immune mROIs, and conversely, immune signatures in IR and bTIL regions. Keratinocyte signatures were enriched in skin adjacent to MIS and IR (**Fig. 6C**), as expected.

Moreover, results were consistent between PickSeq and GeoMX.

To investigate molecular determinants of the spatial heterogeneity within vertical growth phase melanoma revealed by spatial-lag analysis, we performed differential expression (DE) analysis on the IM and EM domains of MEL1; this uncovered 81 significantly upregulated genes in IM and 69 upregulated genes in EM (FDR < 0.05) (**Fig. 6D; Supplementary Table S6**). In IM, GSEA revealed significant enrichment of KRAS signaling and the downstream NF-κB and MYC programs (**Fig. 6E** and **6F**). Upregulation of the KRAS pathway is expected in a tumor such as MEL1 that is mutant in NF1, which functions as a RAS GTPase-activating protein (GAP) (37). BCL2A1 (71), an antiapoptotic pro-survival member of the BCL2 gene family, was expressed in IM but not in EM (**Fig. 6F**). EMT-associated genes were also differently expressed: the S100A4 metalloproteases, β-catenin, and vimentin (DMKN, MMP2, CTNNB1, and VIM genes) were upregulated in IM and GSEA analysis confirmed enrichment of an EMT-associated signature within this region (**Fig. 6G** and **Supplementary Fig. S6B**). EMT-related genes are known to promote invasion and metastasis in many human neoplasms (72), consistent with the observed invasion of this melanoma into the underlying dermis. In contrast, an RNA sensing protein DDX58/RIG-I implicated in the suppression of cancer migration (73) was upregulated in EM (P < 0.05) (**Supplementary Fig. S6C**). The insulin-like growth factor receptor IGF-1R and the IGF binding protein IGFBP2, which is a mitogenic factor (74), were significantly upregulated in EM relative to IM (**Fig. 6F**). Thus, even though IM and EM are contiguous and both in the vertical growth phase, they exhibited significant differences in mitogenic, survival, and EMT-associated programs.

To identify genes differentially expressed with tumor progression, we compared mrSEQ data of tumor in aggregate (EM plus IM) with MIS; this yielded 1,327 DE genes (FDR < 0.05) (**Supplementary Table S6**). However, differences in cellular composition were a complicating factor in this analysis: EM and IM contained mostly tumor cells with very few immune cells, but MIS was rich in immune cells and keratinocytes in addition to tumor cells. To correct for this effect, we searched for a gene shown by imaging and mrSEQ to be expressed in SOX10^+^ tumor cells from EM and IM but not in MIS and then constructed a correlation-based gene network to identify genes co-expressed with that gene (see methods); S100B was found to be an ideal candidate for this purpose (epidermal Langerhans cells also stain positive for S100B, but they were too infrequent to affect the analysis; **Fig. 6H**). The resulting S100B correlation module comprised 35 genes (at r = 0.6) all of which exhibited statistically significant upregulation in EM-IM (FDR < 0.05) (**Fig. 6I** and **6J; Supplementary Table S6**). Among these genes, we validated by CyCIF the upregulation of CD63 and PMEL at the protein level (**Fig. 6K**). The S100B module included: (i) genes implicated in metastasis or invasion in diverse cancers such as SERPINE2 (75), CTSL (76, 77), TBC1D7 (78), and NRP2 (79); (ii) MITF-regulated genes such as the SCD (80) and CDK2 (81); (iii) oncogenes, such as the ETV5 transcription factor (82, 83). When we examined TCGA melanoma data we found that multiple genes in the S100B module (BRI3, CDK2, MT-ND2, PMEL, SOX10, TBC1D7, TSPAN10, TYR) were associated with lower survival (P <0.05) (**Supplementary Fig. S6D**). Thus, half of the genes differentially expressed between MIS and EM-IM have established roles in oncogenesis, invasion, or progression in one or more cancers and ∼25% are associated with lower survival in melanoma. This gene set may warrant further analysis as a means to refine current approaches to determining melanoma risk by gene set analysis (e.g., using methods such as DecisionDx-Melanoma) (84).

### Tumor-immune interaction induces multiple immune suppression programs at the invasive boundary

To better understand invasive properties of primary melanoma (**Fig. 7A**), we combined mrSEQ, conventional CyCIF (with a total of 80 antibodies in five separate panels), and 3D high-resolution deconvolution microscopy from tumor MEL1. In the invasive boundary (IB) region, mrSEQ data revealed significant and localized upregulation of IFNɣ and JAK-STAT signaling as well as the IFN-inducible cytokines *CXCL10* and *CXCL11* (**Fig. 7B** and **Supplementary Fig. S7A-S7C**). *CXCL10* and *CXCL11* (along with *CXCL9* and the *CXCR3* receptor) have diverse roles in regulating immune cell migration, differentiation, and activation, and play a role in response to immune checkpoint inhibitor therapy (85). IFNɣ mediated JAK-STAT signaling can promote upregulation of the metabolic enzyme *IDO1* (86) that has previously been reported to inhibit CTL activation (87, 88) and also promote recruitment of regulatory T cells and myeloid-derived suppressor cells (MDSCs) (89). Consistent with this observation, we observed spatially restricted expression of *IDO1* at the IB (**Fig. 7B**). Additionally, both mrSEQ and CyCIF of tumor cells revealed spatially restricted expression of MHC-II (*HLA-DPB1*) (**Fig. 7C** and **Supplementary Fig. S7D**), which is known to be IFNɣ-inducible (90). MHC-II binds to LAG3 on CD4+/ CD8+ T cells, promoting melanoma persistence by upregulating MAPK/PI3K signaling and can also facilitate immune escape by suppressing FAS-mediated apoptosis (91). Thus, a tightly restricted microenvironment exists at the IB involving multiple cytokines that induce, and are induced by, the JAK-STAT-IDO1 pathway (92) leading to the formation of a highly localized immune-suppressive environment.

**Figure 7:**
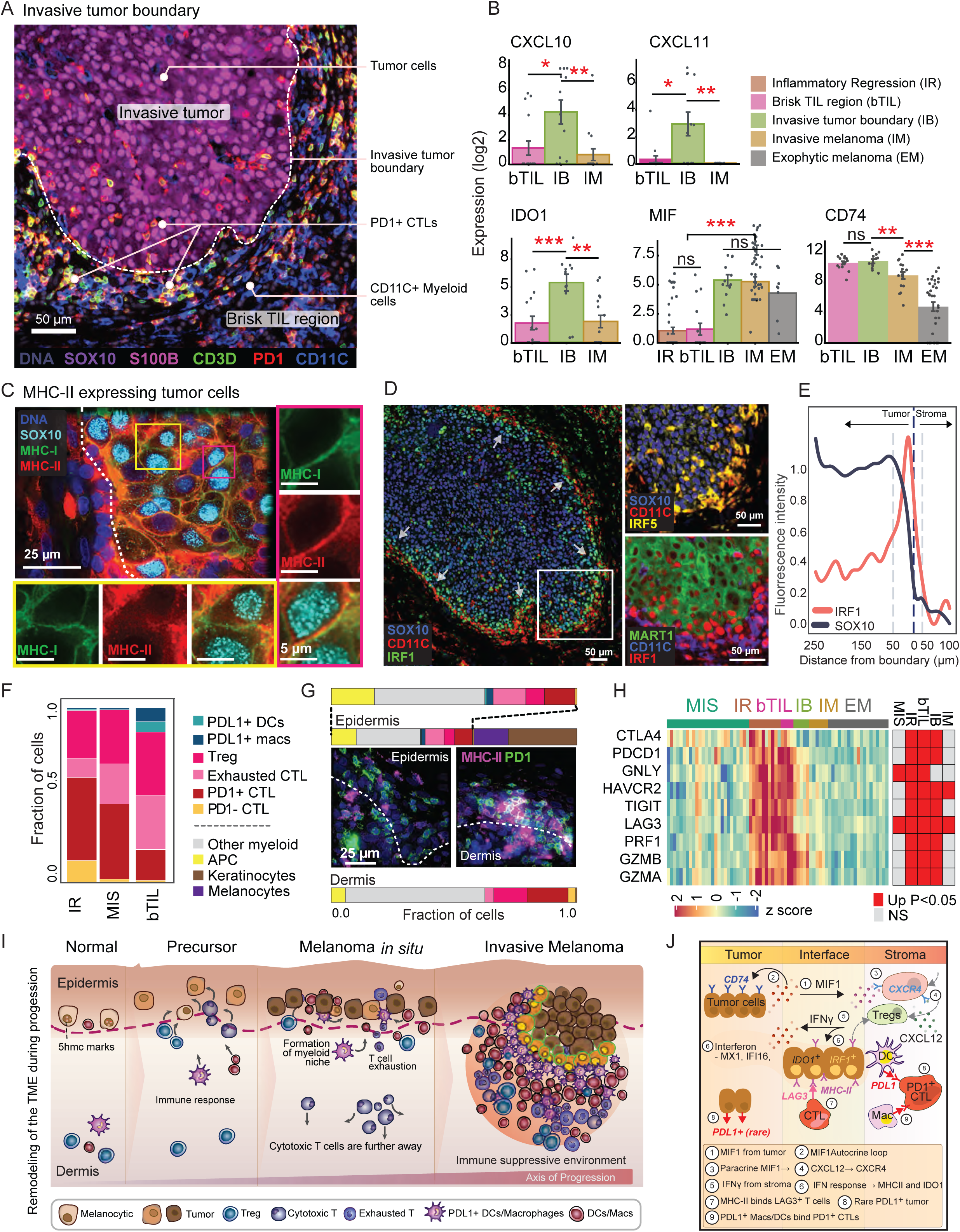
(A) CyCIF image of specimen MEL1-1 showing a protruding edge of the invasive tumor (SOX10: violet, S100B: pink) into the dermis; outside the tumor boundary marked by a white line (dashed) is the brisk TIL region, which contains activated/exhausted T cells (CD3: green, PD1: red) and myeloid cells (CD11C: blue). Scale bar, 50 µm. **(B)** Expression of CXCL10, CXCL11, IDO1, MIF, and CD74 among histological sites (PickSeq data). Values represent mean ± SEM; *P <0.05, **P <0.01, ***P <0.001, ns = not significant. **(C)** CyCIF field of view of MEL1-1 highlighting the spatial arrangement of MHC-II^+^ tumor cells at the invasive front. Tumor cells were stained with SOX10 (cyan), MHC-I (HLA-A: green), and MHC-II (HLADPB1: red). Magnified regions outlined in magenta and yellow squares illustrate MHC-II^+^ and MHC-II^-^ staining of tumor cell membranes. Scale bars, 25 µm (main image) or 5 µm (insets). **(D)** CyCIF of specimen MEL1-1; (left) zoomed out view of invasive front stained for melanocytes (SOX10: blue), myeloid cells (CD11C: red), and interferon signaling (IRF1: green); (right-top) zoomed-in view of invasive front apex stained for melanocytes (SOX10: blue), myeloid cells (CD11C: red) and interferon signaling (IRF5: yellow); (right bottom) zoomed-in view of invasive front apex stained for melanocytes (MART1: green), myeloid cells (CD11C: blue) and interferon signaling (IRF1: red). Scale bar, 50 µm. **(E)** Line plot showing scaled fluorescence intensity of SOX10 (blue) and IRF1 (pink) within (tumor; left of the dashed blue line) and outside (stroma; right of the dashed blue line) the invasive tumor front seen in panel D. **(F)** Stacked bar graph showing the proportions of lymphoid and myeloid cells between the histological regions (IR, MIS, bTIL) in specimen MEL1-1. **(G)** CyCIF maximum-intensity projection images of MEL1-1 of the region of inflammatory regression (shown in panel 1C). Fields are stained for DNA (blue), PD1 (green), and MHC-II (HLA-DPB1: magenta). The dermal-epidermal junction is indicated with a dashed white line. The bar plot shows the proportions of all cell types in the epidermis (upper plots), with lymphocyte and myeloid subset further highlighted, and in the dermis (lower plots); color code is as in panel F. Scale bar 25 µm. **(H)** Heatmap showing expression of genes related to immune checkpoints and T cell activation between histological mROIs in patient MEL1 (GeoMX). Significant upregulation in comparison to the EM region (P <0.05) is highlighted in red, non-significant in grey. **(I)** Schematics of remodeling of the tumor microenvironment with disease progression; see text for details. **(J)** Summary of mechanisms of immune suppression detected in sample MEL1-1.

Expression of other interferon-stimulated genes (ISGs) such as *IRF1* and *IRF5* was also evident at the IB: imaging revealed nuclear staining of IRF1 in tumor cells and strong IRF5 staining in CD11C^+^ myeloid cells directly adjacent to the tumor boundary (**Fig. 7E**, right panels, and **Supplementary Fig. S7E**). By integrating protein intensities across this boundary, we found that the half-maximal width for IRF1 staining was ∼40 µm (**Fig. 7E**) and that of MHC-II expression roughly twice as wide (i.e., ∼ 100 µm or 4 cell diameters; **Fig. 7C** and **Supplementary Fig. S7D**). Thus, mrSEQ and imaging are consistent with a paracrine signaling mechanism in which IFNɣ arising in the peritumoral stroma (including the bTIL region) diffuses into the tumor, inducing ISGs at the invasive front (93).

A reciprocal mechanism involved the macrophage migration inhibitory factor (*MIF*), an inflammatory cytokine overexpressed in a variety of cancers (94). *MIF* was more abundant in tumors (MIS, IM, EM) than in immune-rich regions (bTIL, IR; DE with P <0.05) (**Fig. 7B**) and was confirmed by imaging (**Supplementary Fig. S7F**). mrSEQ data showed that the *MIF* receptor *CD74* (which is induced by IFNɣ (95)), was expressed in immune-rich (bTIL) regions adjacent to the IB (**Fig. 7B**) and CyCIF confirmed this at the protein level (**Supplementary Fig. S7G**). *CD74* was also found to be expressed in melanoma cells (where it can promote PI3K/AKT activation and cell survival) but was spatially restricted to cells at the IB (**Supplementary Fig. S7G**). We also detected elevated expression of a second *MIF* receptor, *CXCR4*, and another cognate ligand, *CXCL12*, in the bTIL region; *CXCR4* activation leads to expansion of immunosuppressive Tregs (96). *CXCR4* is the chemokine receptor most commonly found on cancer cells, and binding to *CXLC12* is thought to promote invasive and migratory phenotypes leading to metastasis (97). However, mrSEQ showed that *CXCR4* levels were low in IM and EM (**Supplementary Fig. S7H**). Thus, mrSEQ data are most consistent with *MIF* expression in tumor cells that acts in a paracrine manner on immune cells with overlapping *CXCL12*-*CXCR4* signaling, also in the immune compartment. Overall, these data reveal the pattern of immune cell activation and immunosuppression involving highly localized cytokine signaling and direct cell-to-cell contact all within a few cell diameters of the IB.

However, successful immune editing and clearance of SOX10^+^ tumor cells were also observed at regions of inflammatory and terminal regression in MEL1; only a few millimeters away from the invasive tumor front. In regions of regression, we observed dense infiltrates of CTLs, the majority of which were PD1^+^ and thus activated. The greatest concentration of PD1^-^ CTLs in MEL1 was found not near the tumor but in the IR region (**Fig. 7F** and **Supplementary Fig. S7I**). MHC-II^+^ APCs were also abundant, consistent with ongoing Treg activation (**Fig. 7G** and **Supplementary Fig. S7I**). Imaging showed that CTLs in the IR that were PD1^+^ also expressed LAG3 and/or TIM3 and mrSEQ confirmed the expression of PDL1, LAG3, TIGIT, and CTLA4. Thus, many T cells in the region of IR appeared to be exhausted. An accumulation of terminally exhausted T cells near the tumor is likely to reflect normal immune homeostasis (immune induced tumor regression), not tumor-mediated immune suppression (**Fig. 7H**). Our data suggest that the key difference between the active immune response in regions of tumor regression and the seemingly ineffective response in regions of invasion is the presence of abundant PDL1^+^ macrophages and dendritic cells and their physical interaction with T cells.

## DISCUSSION

In this paper, we exploit histological features used clinically to stage primary cutaneous melanoma as a framework for analyzing multiplexed imaging and mrSEQ data along an axis of tumor progression from precursor fields to invasive melanoma. We also examine regions near the dermal-epidermal junction in which immunoediting is ongoing or had reached a resolution with few or no tumor cells remaining.

Molecular evidence of progression was obtained using protein markers (by CyCIF) and oncogenic programs (by mRNA expression) both within specimens, each of which comprised several distinct histologies, and also across a patient cohort. Disease-relevant morphological features ranged in length scale from 0.5 µm (organelles) to 20 mm (invasive fronts) and we found that imaging the entirety of individual specimens up to ∼1 cm in length – not a TMA or a small region of interest – was essential for retaining information on tissue context and for the success of our approach (34). Accompanying high-resolution 3D imaging revealed the presence of immune synapses and PD1-PDL1 co-localization to the plasma membranes of neighboring cells; we interpret these as evidence of functional cell-to-cell interaction. Juxtacrine receptor-ligand interactions of this type appear to be relatively common among cells lying along the immune-rich invasive tumor boundary (up to 20% of adjacent cells making contact in an exemplary field). At the current state of the art, however, only a relatively small number of whole-slide images could be analyzed for spatial patterns in their entirety (n = 13 patients and 70 histological ROIs). Thus, the progression-associated changes described in this manuscript should be regarded as representative rather than comprehensive: in contrast, discovery of new (progression) biomarkers by traditional IHC typically involves analysis of at least 100 specimens followed by clinical trials (98).

The use of Latent Dirichlet Allocation (LDA) on high-plex data made it possible to identify recurrent combinations and arrangements of cell types across ROIs. The frequency of recurrent cellular neighborhoods (RCNs), and their proximity to each other, changed with disease stage (**Fig. 7I**). Relative to adjacent normal skin, changes in the immune environment were detectable in fields of melanocytic atypia (precursor fields) but the largest differences along the progression axis appeared to involve precursor fields and MIS. This involved the recruitment to the tumor domain of CTLs, many of which were PD1^+^ and presumably activated as well as increases in suppressive Tregs and PDL1-expressing myeloid cells. The resulting immunosuppressive environment became more consolidated between MIS and invasive stages. For example, in sample MEL1, a community of cells involving tumor and PDL1^+^ myeloid cells (macrophages and dendritic cells in roughly equal proportion) formed a thin and continuous sheath along the invasive front. TILs were largely excluded from the tumor at this stage, except in the immediate proximity of small vascular structures that were found throughout the EM.

Whereas LDA was effective at identifying neighborhoods involving different types of cells, spatial lag modeling on CyCIF data identified recurrent patterns involving continuous differences in protein levels on a scale of 10 to 100 cell diameters. Spatial gradients on similar scales were also observed for several protein markers – MITF or S100B for example. Thus, whereas LDA and clustering of transcriptional data highlight discrete differences in cell states, imaging demonstrates the presence of gradients reminiscent of those found in developing tissues (60, 61). In general, significant differences among communities of cancer cells identified by shift-lag modeling involved hyperdimensional features (combinations of markers instead of single proteins) consistent with the current understanding of molecular determinants of cellular morphology (99). Moreover, gradients in MITF or S100B are likely to be indications that tumor cell phenotypes are graded in space but not causes of this variation. One unexpected finding involved the “invasive” state of melanoma cells, which is often described as being MITF^low^ with slow proliferation. Spatial lag modeling showed that MITF^high^ KI67^high^ cells were common in MEL1 in the immediate proximity of the invasive front and mrSEQ showed that these cells were significantly enriched in EMT-associated programs, which are common along the invasive boundaries of many other types of tumors. Future studies on paired primary and metastatic tumors will be required understand how these data relate to previous analysis of MITF high and low states, which has largely been performed in cell lines.

CyCIF and mrSEQ revealed a pattern of cytokine production and receptor expression at the invasive boundary of MEL1 consistent with paracrine regulation of both tumor and immune cells (**Fig. 7J**). The dermis in this region was rich in TILs (corresponding to a brisk TIL response in the Clark grading system) and was the site of highest IFNɣ production. A band of cells ∼2 cell diameters wide in the adjacent invasive melanoma stained positive for nuclear-localized IRF1, the master regulator of interferon response (yellow cells in **Fig. 7I**); mrSEQ showed that JAK-STAT signaling was active in this region and IDO1 was differentially expressed. IDO1 converts tryptophan into kynurenine, which activates Tregs and MDSCs, and is known to be immunosuppressive in melanoma (100). MHC-II was also expressed in both immune and tumor cells at the invasive boundary, in a band roughly twice as wide as IRF1, and is known to function in this context by binding to LAG3 on TILs, leading to inhibition of TCR signaling and T cell activation (101, 102). MIF1 was another inflammatory cytokine found at the invasive front and was expressed primarily in the invasive tumor region; responsive CXCR4-expressing immune cells were found in the stroma. MIF1 may also have an autocrine activity since expression of the MIF1 receptor CD74 was detected in the tumor itself. CXCR4-expressing immune cells are also responsive to CXCL12, which was expressed in the TIL-rich stroma. CXCR4 is the cytokine receptor most commonly found on melanoma and other types of cancer cells, and CXCR4- CXCL12 signaling is thought to promote metastasis (97), but we did not observe CXCR4 expression in MEL1 by mrSEQ. We conclude that the immunosuppressive activity of IFNɣ manifests itself in MEL1 in a spatially restricted manner involving a sheath of tumor and myeloid cells surrounding the invasive tumor. Undoubtedly, the magnitude of these effects will vary among primary tumors but our analysis illustrates how reciprocal cytokine signaling between tumor and immune cells can shape the local TME.

Performing spatial proximity analysis on imaging data (with a 10 - 20 µm cutoff) made it possible to identify cells that are sufficiently close to each other that physical contact is probable. We were able to visualize these contacts and infer function using high-resolution 3D imaging of ∼5 x10^3^ cells. The most informative images were those involving cytotoxic T and melanoma cells that resulted in the polarization of CD8 (a co-receptor for the T-cell receptor) at the point of contact, suggesting the formation of a synapse. PD1^+^ CTLs cells were also observed in contact with PDL1-expressing macrophages and dendritic cells resulting in receptor-ligand co-localization. In some cases, these contacts involved surprisingly extended processes (>10 µm) in which macrophages appeared to stretch towards T cells. In other cases, multiple CTLs, T helper, and myeloid cells were found to be in physical contact with each other and with tumor cells with evidence of receptor or ligand polarization. The functional significance of these clusters awaits further analysis using a greater diversity of immune markers, but they are presumably a physical manifestation of the competing activating and inhibitory effects of other immune cells on CTLs.

Overall, we found evidence of at least six immunosuppressive mechanisms operating near the invasive front. Particularly striking was the overlap in the binding of PD1^+^CTLs to PDL1^+^ macrophages and dendritic cells and tumor cell-intrinsic phenotypes such as MHC-II and IDO1 expression. Unexpectedly we only rarely detected high expression of PDL1 on tumor cells by either whole-slide imaging or high-resolution microscopy (even when IFNɣ expression was detected). This finding was validated using a separate cohort of 25 primary cutaneous melanomas and contrasts with data collected in parallel from metastases, in which strongly PDL1^+^ tumor cells are common. We conclude that myeloid cells are likely to provide the predominant source of PDL1 bound to PD1^+^ T cells in the tumors in our cohort. Data obtained by Oh et al. (52) in mouse models suggest that the functionally significant cell type is likely to be PDL1^+^ dendritic cells, but in our specimens, dendritic cells and macrophages appear to be similar in abundance at tumor invasive boundaries.

CTLs were found to engage tumor cells in a region of inflammatory regression adjacent to MIS in MEL1. The additional presence of an adjacent region of complete regression, which was rich in immune cells but free of tumor cells, suggests that immune editing was ongoing and successful. However, these regions also had a preponderance of terminally exhausted CTLs, showing that the characteristics of a successful and self-limiting anti-tumor immune response in data such as that presented here can resemble those of immunosuppression in invasive melanoma. The primary difference we observed between regions of regression and invasion with immunosuppression was a substantially lower level of PDL1^+^ myeloid cells, but further research will be required to determine if this is generally true.

## Limitations of this study

One challenge encountered in molecular analysis of primary melanoma is that, as a diagnostic necessity, specimens are available only in FFPE form, generally precluding scRNA-Seq for research purposes.

Sequencing of carefully selected micro-regions by mrSEQ provides meaningful information on activated pathways and differential gene expression linked to histological features but is not yet single-cell resolution. A second challenge is that meaningful outcome analysis requires long follow-up: all patients whose tumors were analyzed in this study were diagnosed between 2017 and 2019 and were alive at the time of the last follow-up; ∼75% were disease-free. Thus, we used histologic progression not outcome to organize the data in a biologically meaningful fashion. A final limitation in any molecular study of patient-derived specimens is that only one-time point can be evaluated per patient. Our analysis of tumor samples exhibiting progression within the same specimen helps to mitigate this issue.

Despite the scope of the current data collection effort, 13 specimens are too few to be representative of the diversity of cutaneous melanoma. We estimate that data collection will need to be scaled up 5 to 10- fold to determine whether many of the features observed in MEL1 are significantly associated with progression in other specimens. Moreover, spatially resolved mRNA expression and high-plex imaging data support each other in many cases, but this was not always true. This is not unexpected because mRNA and protein expression are known to be uncorrelated in many cases (103) and cell morphology represents a hyper-dimensional feature in gene expression space (99). 3D image data has provided valuable insight into cell-to-cell interactions, but automated segmentation of these data remains difficult, and most conclusions were derived from a human inspection of images. More generally, the greatest limitation in the current work is related to the underdevelopment of software tools for characterizing large high-plex tissue images. Much, therefore, remains to be discovered from the images we have collected. Full resolution Level 3 images (104) and associated single-cell data are therefore being released in their entirety, without restriction, for follow-on analysis.

## METHODS

### Contact for reagent and resource sharing

This manuscript does not contain any unique resources and reagents; all data is provided for download without restrictions. Any questions should be directed to the lead contact Peter Sorger.

### Clinical samples

Using medical records and pathological review of hematoxylin and eosin (H&E) stained diagnostic specimens, we retrospectively identified 13 patients with tissue samples containing various stages of melanoma progression (**Supplementary Table S1** and **S2**). The samples were retrieved from the archives of the Department of Pathology at Brigham and Women’s Hospital and collected under the Institutional Review Board approval (FWA00007071, Protocol IRB18-1363), under a waiver of consent. Fresh formalin-fixed paraffin-embedded (FFPE) tissue sections were cut from each tumor block. The first section of each block was H&E stained and used to annotate regions of interest (ROIs; **Supplementary Table S3**). The remaining subsequent FFPE slides were used for cyclic multiplex immunofluorescence imaging (CyCIF) experiments to characterize markers of melanoma progression and the features of the immune microenvironment within various stages of melanoma. A specimen from a single patient MEL1 (samples MEL1-1, MEL1-2, and MEL1-3) was selected for deeper profiling with CyCIF and high-resolution imaging, in addition to microregion transcriptomics (PickSeq, GeoMX). The clinical, biospecimen, and imaging level metadata were all collected following the MITI standards (104).

Based on the melanoma diagnostic criteria, the histopathological annotations included normal skin (N), melanoma precursor lesions (P: melanocytic atypia, dysplasia, and hyperplasia), melanoma in situ (MIS), vertical growth phase of melanoma (VGP), radial growth phase of melanoma (RGP), invasive (IM) and nodular melanoma (NM); the exophytic component of the polypoid melanoma was labeled as exophytic melanoma (EM). These ROIs were further classified and subdivided based on the presence of immune infiltrate (brisk TIL (bTIL), inflammatory regression (IR), none) and various histologically distinct structures (epidermis, dermis, invasive front (IB)). The bTIL region was defined as a dense lymphocytic infiltrate in the stroma adjacent to the invasive tumor. IB was defined as the tumor region extending ∼20 μm from the tumor-stroma interface. The most representative regions of each histologic category from each specimen were selected in order to avoid inter-observer variability. In the case that a single specimen contained more than one histologic region in each category (e.g., precursor regions on both sides of VGP melanoma), we performed neighborhood analyses separately since these regions were not physically adjacent.

### Imaging (H&E and t-CyCIF)

H&E stained FFPE slides were digitized using an Olympus VS-120 automated microscope using a 20x objective (0.75 NA) at the Neurobiology Imaging core at Harvard Medical School. CyCIF was performed as described in (31) and at protocols.io (dx.doi.org/10.17504/protocols.io.bjiukkew). In brief, the BOND RX Automated IHC Stainer was used to bake FFPE slides at 60°C for 30 min, dewax using Bond Dewax solution at 72°C, and perform antigen retrieval using Epitope Retrieval 1 (Leica^TM^) solution at 100°C for 20 min. Slides underwent multiple cycles of antibody incubation, imaging, and fluorophore inactivation. Antibodies were incubated overnight at 4°C in the dark; in contrast to the protocol.io method, this was performed using a solution that also included Hoechst 33342 for DNA staining. Before imaging, glass coverslips were wet-mounted using 100 μL of 70% glycerol in 1x PBS. Images were acquired using a CyteFinder® slide scanning fluorescence microscope (RareCyte Inc., Seattle WA) with a 20x/0.75 NA objective. Slides were soaked in 42°C PBS to facilitate coverslip removal; then fluorophores were inactivated by incubating slides in a solution of 4.5% H2O2 and 24 mM NaOH in PBS and placing them under an LED light source for 1 hr. The list of all antibody panels used in the experiments is presented in **Supplementary Table S4**. All the used antibodies were validated with a multi-step process including comparing multiple antibodies with each other and with clinical standards, and by visual inspection on individual stained FFPE tissue sections. Antibodies that passed these criteria and followed the expected staining pattern were only included in downstream anslysis.

One FFPE section from sample MEL1-1 was imaged with CyCIF at high-resolution using a DeltaVision ELITE microscope (Cytiva; formerly GE Sciences) equipped with a 60x/1.42NA oil-immersion objective and an Edge 4.2 (PCO) sCMOS camera. For accurate deconvolution, an oil refractive index of

1.524 was selected through optimizing multiple acquired point-spread functions as it provided the highest image quality. The slide was wet-mounted with a high-precision 1.5-grade coverslip (ThorLabs CG15KH1) using 105 μL of 90% glycerol. The fields for image acquisition were selected by evaluating SOX10 staining to locate and identify melanocytes and tumor cells, yielding a total of 42 fields across the annotated regions (**Fig. 1C** and **Supplementary Fig. S1D**). Images were acquired in 5 μm Z-stacks at 200 nm step size to create a 3D representation of the sample. Excitation wavelengths were: 632/22 nm, 542/27 nm, 475/28 nm, 390/18 nm for four-channel imaging.

PD-L1 expression was also quantified in an additional cohort of 25 additional primary melanomas (cohort 2; **Fig. 4F** and **G**) selected from the BWH tissue bank using the same criteria as the 13 specimens described above. These specimens were subjected to lower-plex CyCIF imaging using antibodies listed in **Supplementary Table 4**. The frequency of PDL1 positivity on tumor and myeloid cells was then visually quantified as the percentage of PDL1-positive SOX10^+^ tumor cells (binned as follow: 0-5%, 5-20%, >20%) or CD11C^+^ myeloid cells (binned as follows: <1%, 1-25%, >25%). Broad bins were chosen to make the results robust to counting errors in regions of tissue where cells were densely packed.

### Microregion transcriptomics

For the microregion transcriptomic profiling (mrSEQ) using PickSeq and GeoMX, we identified micro- regions (mROIs) of MIS, EM, IM, IB, IR, and bTIL from samples MEL1-1, -2, and -3 based on the corresponding H&E-stained sections. Freshly cut serial sections from the corresponding tissue blocks were used for the mrSEQ experiments.

### PickSeq processing and library preparation

PickSeq is a method by which 40 µm mROIs of interest are physically extracted using a robotic arm followed by mRNA extraction and RNA sequencing (32). 222 ROIs representing five morphologically distinct sites (MIS, IM, IB, bTIL, EM; **Supplementary Fig. S6A**) were selected for collection and library preparation. The FFPE sections were deparaffinized and rehydrated using the Histogene Refill Kit (Arcturus). Slides were immersed in xylene for 5 min, a second jar of xylene for 5 min then incubated in a series of ice-cold solutions with 0.0025% RNasin Plus (Promega): 100% ethanol for 1 min, 95% ethanol for 1 min, 75% ethanol for 1 min, 1X PBS for 1 min, and another tube of 1X PBS for 1 min. Slides were stained with 50 µM DRAQ5™ a Far-Red DNA Dye (ThermoFisher) in PBS, with 0.1% RNasin Plus for 2 min on ice. Sections were dehydrated in a series of ice-cold solutions with 0.0025% RNasin Plus: 1X PBS for 1 min, 1X PBS for 1 min, 75% ethanol for 1 min, 95% ethanol for 1 min, 100% ethanol for 1 min. Slides were left in ice-cold 100% ethanol before mROI retrieval.

For mROI retrieval, the slides were loaded into a CyteFinder instrument (RareCyte) and retrieved using the integrated CytePicker module with 40 µm diameter needles. The retrieved tissue mROIs were deposited with 2 µl PBS into PCR tubes containing 18 µl of lysis buffer: 1:16 mix of Proteinase K solution (QIAGEN) in PKD buffer (QIAGEN), with 0.1% RNasin Plus. After deposit, tubes were immediately placed in dry ice and stored at -80°C until ready for downstream RNA sequencing workflow.

PCR tubes containing tissue microregions in the lysis buffer were removed from the freezer, allowed to thaw at room temperature for 5 min, and incubated at 56°C for 1 hr. Tubes were briefly vortexed, spun down, and placed on ice. Dynabeads Oligo(dT)25 beads (ThermoFisher) were washed three times with ice-cold 1X hybridization buffer (NorthernMax buffer (ThermoFisher) with 0.05% Tween 20 and 0.0025% RNasin Plus) and resuspended in original bead volume with ice-cold 2x hybridization buffer (NorthernMax buffer with 0.1% Tween 20 and 0.005% RNasin Plus). A volume of 20 µl of washed beads was added to each lysed sample, mixed by pipette, and incubated at 56°C for 1 min followed by room temperature incubation for 10 min. Samples were placed on a magnet and washed twice with an ice-cold 1X hybridization buffer, then once with ice-cold 1X PBS with 0.0025% RNasin Plus. The supernatant was removed, and the pellet was resuspended in 10.5 µl nuclease-free water. Samples were incubated at 80°C for 2 min and immediately placed on a magnet. The supernatant was transferred to new PCR tubes or plates, and placed on ice for subsequent whole transcriptome amplification or stored at -80°C.

Reverse transcription and cDNA amplification were performed using the SMART-Seq v4 Ultra Low Input RNA Kit for Sequencing (Takara Bio, Kusatsu, Shiga, Japan). The resulting amplified cDNA libraries were assessed for DNA concentration using the Qubit dsDNA HS Assay Kit (ThermoFisher) and for fragment size distribution using the BioAnalyzer 2100 High Sensitivity DNA Kit (Agilent). The sequencing libraries were prepared with ThruPLEX DNA-seq Kit (Takara Bio). The resulting libraries were characterized by using the Qubit dsDNA HS Assay Kit and BioAnalyzer 2100 High Sensitivity DNA Kit, pooled at equimolar ratios, and sequenced using a MiSeq (Illumina) or NextSeq (Illumina) sequencer.

### GeoMX processing and data collection

NanoString GeoMx gene expression analysis utilizing the cancer transcriptome array (CTA) probe set was performed by the Technology Access Program at NanoString using previously described methods (33). Briefly, a 5 μm section of FFPE melanoma was dewaxed and stained overnight with antibodies targeting melanocytes (PMEL), epithelial (pan-cytokeratin), and immune cells (CD45) defining cell morphology and highlighting regions of interest. The section was hybridized with the CTA probes before being loaded into the instrument. Seventy ROIs representing five morphologically distinct sites (MIS, IM, IB, bTIL, EM; **Supplementary Fig. S6A**) were selected for collection and library preparation. All sample processing and sequencing were performed by the Technology Access Program at NanoString. Probe measurements, and quality control data were provided by NanoString.

### 3D image processing, alignment, and visualization

Acquired images were deconvolved using constrained iterative in SoftWorx to reassign photons to the focal plane and increase image contrast. Maximum intensity projections were also generated.

Subsequently, cycles were aligned using a custom script written in MATLAB (Mathworks). Briefly, 2D image registration was first carried out using the Hoechst channel maximum intensity projections. This was followed by registration along the z-axis. The registered 3D datasets were visualized in Imaris (Bitplane) and surface rendered for visualization.

### PickSeq data Alignment and expression matrix generation

The raw FASTQ files were examined for quality issues using FastQC (http://www.bioinformatics.babraham.ac.uk/projects/fastqc/) to ensure library generation and sequencing were suitable for further analysis. The reads were processed using the *bcbio* pipeline v.1.2.1 software (105). Briefly, reads were mapped to the GRCh38 human reference genome using *HISAT2* and Salmon. Length scaled transcripts per million (TPM) derived from *Salmon* were used for the downstream analysis.

### Differential gene expression and pathway analysis

*DESeq2* R package was used to generate the normalized read count table based on their estimateSizeFactors() function with default parameters by calculating a pseudo-reference sample of the geometric means for each gene across all samples and then using the "median ratio" of each sample to the pseudo-reference as the sizeFactor for that sample. The sizeFactor was then applied to each gene’s raw count to get the normalized count for that gene. *DESeq2* (106) was used for differential gene expression analysis. A corrected P-value cut-off of 0.05 was used to assess significant genes that were up-regulated or down-regulated using Benjamini-Hochberg (BH) method. Principal component analysis (PCA) was performed using the *prcomp* R package. A compendium of biological and immunological signatures was identified from publicly available databases or published manuscripts for performing enrichment analysis. To perform gene set enrichment analysis, two previously published methods (Gene Set Enrichment Analysis (GSEA) and single-sample GSEA (ssGSEA)) were primarily used. The R package *clusterProfiler* was used to perform GSEA and the R package *GSVA* was used to perform ssGSEA which calculates the degree to which the genes in a particular gene set are coordinately up- or down-regulated within a sample. The KRAS and JAK-STAT were curated from MSigDB (107), and immune cell-related and melanoma-related signatures were curated from published studies (7,108,109).

### Network analysis to identify genes within S100B module

The normalized expression matrix (PickSeq data) was loaded into the network analysis tool BioLayout (110). Within the tool, a Pearson correlation matrix was generated, i.e., an all versus all comparison of expression profiles across all samples. A gene correlation network (GCN) was then generated using a correlation threshold value 0.6. In the context of a GCN, nodes represent genes and edges represent the correlations between them. A single-step neighbor walk was performed within the tool from S100B to determine the S100B module.

### CyCIF image preprocessing and quality control

The complete preanalytical CyCIF image processing (stitching, registration, illumination correction, segmentation, and single-cell feature extraction) was performed using the MCMICRO pipeline (36) an open-source multiple-choice microscopy pipeline, versions 60929d5b82 and 7547d0c42a (full codes available on GitHub https://github.com/labsyspharm/mcmicro). For the generation of probability maps and the nuclei segmentation, a trained U-Net model UnMicst v1 was used followed by a marker-controlled watershed used for single-cell segmentation (111). A diameter range of 3 to 60 pixels was used for nuclei detection. The cytoplasmic area was captured by expanding the nuclei mask by 3 pixels. After generating the segmentation masks, the mean fluorescence intensities of each marker for each cell were computed, resulting in a single-cell data table for each acquired whole-slide CyCIF image. The X/Y coordinates of annotated histologic regions on the whole-slide image were used to extract the quantified single-cell data of cells that lie within the ROI range.

Multiple approaches were taken to ensure the quality of the single-cell data. At the image level, the cross-cycle image registration and tissue integrity were reviewed; regions that were poorly registered or contained severely deformed tissues and artifacts were identified, and cells inside those regions were excluded. Antibodies that gave low confidence staining patterns by visual evaluation were excluded from the analyses. The quality of the segmentation was assessed and the segmentation parameters were iteratively modified to improve the accuracy of the segmentation masks. On the single-cell data level, correlations of DNA staining intensities in different cycles were used to filter out cells that were lost in the cyclic process with a threshold of correlation coefficient less than 0.8.

### Single-cell phenotyping

We first applied unsupervised graph-based clustering approaches such as Leiden and Phenograph on the derived single-cell data (data not shown) to identify cell types discernable in the CyCIF dataset.

However, unlike single-cell RNASeq data, many cell types (especially immune cells) do not form distinct clusters (likely owing to low dimensionality) leading to ambiguity in cell type assignment especially for cells that lay at the boundaries between clusters. Therefore, we developed a gating-based phenotyping approach to classify cells (112). First, an open-source OpenSeadragon based visual gating tool (https://github.com/labsyspharm/cycif_viewer) was used to derive gates (the cut-off value that distinguishes cells that express and do not express a particular marker). The identified gates for each marker were subsequently used to rescale (similar to batch correction) the single-cell data between 0 and 1 such that the values above 0.5 identify cells that express the marker and vice-versa (*rescale* function within *scimap*). We repeated this process on every image independently and merged them into a single large single-cell dataset. The scaled single-cell data was used for cell-type calls.

We built an algorithm (*phenotype_cells* function within the *scimap* python package) that assigns phenotype labels to individual cells based on a sequential probability classification approach. The underlying assumption is that the probability of a real signal would be higher than the bleed-through/ artifact signals that arise due to chromatic or segmentation artifacts. For example, if a B cell (CD20+) and T cell (CD3+) are physically next to each other with some bleed-through of CD3 signal into the B cell (CD20+ cell), the algorithm compares the scaled intensity of CD3 and CD20 and assigns the cell as a B cell due to higher levels of CD20 expression. It is sequential as we follow a tree structure, whereby the cells are initially classified into large groups such as tumor (e.g., based on SOX10/S100B) and immune (CD45 expression) and as a second step, the immune cells are further divided into cell-types such as T cells, B cells, etc. which are further divided into finer subtypes in a sequential step. An input to this algorithm is a relationship chart (phenotyping workflow, **Supplementary Fig. S1C**, **S2B,** and **Supplementary Table S5**) between markers and cell types (phenotypes). Each cell is binned into a phenotype class based on the highest expression of a given marker. If a cell does not express any of the markers (i.e., < 0.5) in the phenotyping workflow sheet, it is assigned to an unknown class. On average we found that ∼25% of cells (15% to 39% across all 13 patients) were classified as unknown. Based on inspection of H&E images these cells are likely to include fibroblasts, adipocytes, muscle, and other stromal cells. By using “AND, OR, ANY, ALL” as parameters, in combination with “POS or NEG” expression patterns, we were able to define the desired cell types identified via unsupervised clustering and manual inspection of the images. The assigned cell types were then verified by overlaying the phenotypes onto the image using Napari (*image_viewer* function within *scimap*). In total, we assigned phenotype labels to 1.7*10^6^ single cells from 70 CyCIF ROIs corresponding to all progression stages (specimens MEL2-MEL13) and a whole slide dataset from specimen MEL1-1.

### Phenotype co-occurrence analysis

For each cell in the CyCIF dataset, its local neighborhood was captured by querying a radius of 20 µm from the cell centroid as measured by Euclidean distance between X/Y coordinates. The phenotypes of these cellular neighbors were mapped to generate a neighborhood matrix containing the neighbor phenotype for every cell. We then randomly permutated (1,000 times) the neighborhood phenotypes without changing the number of neighbors (to maintain the tissue structure) and generated 1,000 random cell-cell neighborhood matrices. The frequency of all cell-to-cell pairwise proximity from the real neighborhood matrix was compared with the 1,000 randomly generated neighborhood matrices to identify significant proximity or avoidance between pairs of cell types. The p-values were derived for every pairwise proximity according to the following formulas:

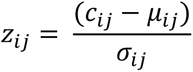

*cij* is the number of times the *ith* cell type was found proximal to the *jth* cell type. Its associated P-value *pij* was calculated by

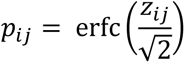

where erfc is the complementary error function calculated using the python function ‘*scipy.stats.norm.s*f’. The method is implemented under the *spatial_interaction* function in the *scimap* python package.

### Spatial lag analysis to define tumor cell communities

For each tumor cell in the CyCIF dataset (MEL1-1), its local neighborhood was captured by querying a radius of 20µm from the center cell as measured by Euclidean distance between X/Y coordinates. A spatial lag vector was derived for each neighborhood by taking the product of the expression matrix and a weighted proximity matrix. The weights were assigned such that the closest cell within a neighborhood received the highest weight (weight = 1) and the farthest received the lowest weight (weight = 0). The weights were then normalized to account for the number of cells within each neighborhood. The spatial lag matrix was then clustered using Python’s *scikit-learn* implementation of *KMeans* with k = 20 and manually grouped (hierarchical clustering assisted) into meta-clusters (10 clusters) based on similar expression patterns visualized using a heatmap. The method is implemented under the *spatial_expression* function in *scimap* python package.

### Proximity volume scoring

To quantify the abundance of cell-to-cell proximity between cell types of interest (COI) observed in CyCIF images, we developed a scoring system that weighs user-defined proximity patterns. The proximity volume score is defined as the proportion of COI found in proximity to each other (10 µm) compared to the total number of cells within that image. We calculated the spatial volume score between cell types of interest (tumor and CD11C^+^ myeloid cells) for each image and averaged them across images belonging to the same stage. The scoring is implemented under the *spatial_pscore* function in *scimap* python package.

### Recurrent cellular neighborhood (RCN) analysis to identify microenvironmental communities

For every single cell from specimens MEL1 to MEL13, its local neighborhood was captured by querying a radius of 20µm from the center cell as measured by Euclidean distance between X/Y coordinates. The cells within each neighborhood were mapped to the cell-type assignment made and their frequency within each neighborhood was computed. The frequency matrix was then used for microenvironment modeling using a method called Latent Dirichlet Allocation (LDA) which is commonly used in the natural language processing (NLP) and information retrieval (IR) community.

Python’s *gensim* (https://pypi.org/project/gensim/) implementation of LDA model estimation was used to train the algorithm. The number of latent motifs to be extracted from the training corpus was determined empirically (motifs = 10). The latent vectors (weights) were recovered from the model and clustered using *scikit-learn* implementation of *KMeans* with k = 30. The optimal number of *KMeans* clustering was determined by looking for the elbow point in the computed cluster heterogeneity during convergence (**Supplementary Fig. S2E**). A fairly lenient elbow point (k = 30) was used to capture the maximal variance in our dataset and to account for smaller communities. The clusters were then manually grouped (hierarchical clustering assisted) into meta-clusters (11 clusters) based on similar microenvironmental community patterns. To validate the RCN assignment, these meta-clusters were overlaid on the original tissue H&E-stained and fluorescent images. For example, RCN1 generally mapped to the epidermis capturing structural components of the data whereas RCN8 mapped to regions of immune suppression (with a high abundance of PD1^+^ T cells) capturing communities of functional importance. In parallel, we also derived RCNs using an alternative approach, whereby we directly cluster the cell-type frequency table generated before feeding into the LDA model. We were able to identify similar communities (**Supplementary Fig. S2D**) thereby validating the communities that we describe using an alternative approach. However, we believe the LDA model was more robust to noise compared to directly clustering the cell-type frequency table. The method is implemented under the *spatial_count* function and the LDA approach is implemented under the *spatial_lda* function in *scimap* python package.

### Statistical tests

All statistical tests to infer P-value for significant differences (P < 0.05) in mean were performed using Python’s *scipy* implementation of the t-test.

### Data and software availability

Micro-region sequencing (mrSEQ) data is available via GEO (GSE171888). All full resolution images derived from image data (e.g., segmentation masks) and all cell count tables are available via the NCI- Human Tumor Atlas Network data portal (https://data.humantumoratlas.org/). These data are also available via the Harvard Tissue Atlas Portal. Note that individual files are ∼100GB in size so an AWS S3-compatible download tool should be used, connecting to the following address: s3://hta-melatlas- 1/data/. Exploration is possible via:

- https://hta-melatlas-1.s3.amazonaws.com/data/index.html
- https://labsyspharm.github.io/HTA-MELATLAS-1/

Several of the figure panels in this paper are available with text and audio narration for anonymous online browsing using MINERVA software (113, 114), which supports zoom, pan, and selection actions without requiring the installation of software.

- HTA MEL Atlas 1: Introduction to the MEL Atlas.

## DECLARATION OF INTERESTS

PKS is a member of the SAB or Board of Directors of Glencoe Software, Applied Biomath, and RareCyte Inc. and has equity in these companies; PKS is also a member of the SAB of NanoString and a consultant for Montai Health and Merck. Glencoe, RareCyte, and NanoString provided commercially- available technology used in this study. SS is a consultant for RareCyte Inc. ZM is a consultant for Verseau Therapeutics Inc. The authors declare that none of these relationships have influenced the content of this manuscript.

## Financial support

This work was supported by NIH grants U2C-CA233262 (PKS, SS), K99- CA256497 (AJN), the Ludwig Center at Harvard (PKS, SS), R50-CA252138 (ZM), and by grants from the Finnish Medical Foundation and the Relander Foundation (TV). Access to the GeoMX mrSEQ platform was kindly provided by NanoString Inc. as part of their Technology Access Program. (TAP). All HTAN consortium members are named at (humantumoratlas.org). We thank Dana-Farber/Harvard Cancer Center for the use of the Specialized Histopathology Core, which provided histopathology services supported by P30-CA06516. Imaging at the HMS Neurobiology Imaging Facility (of H&E specimens) was supported by NINDS Core Center Grant P30-NS072030.

## TABLE LEGENDS

**Supplementary Table S1.** Patient characteristics.

**Supplementary Table S2.** Clinical data related to the analyzed specimens.

**Supplementary Table S3.** Distribution of Regions of Interest (ROI) by histopathological annotation and patient ID.

**Supplementary Table S4.** Distribution of the clinical samples MEL1-MEL13 used in each CyCIF experiment (experiments 1 to 5) and the corresponding antibody panels.

**Supplementary Table S5.** The criteria for identifying individual cell phenotypes in CyCIF experiments 1 and 2 (antibody panels presented in Supplementary Table S4).

**Supplementary Table S6**. Gene and protein symbols and names and the differentially expressed genes between MIS and EM-IM or between EM and IM regions.

**Supplementary Table S7**. Mean distance between all RCN’s and RCN10 (tumor enriched) grouped by progression stages.

**Supplementary Video 1**. A synapse between a PDL1+ macrophage and a PD1+ CD8+ T cell.

**Supplementary Minerva story 1.** HTA MEL Atlas 1: Introduction to the MEL Atlas (https://labsyspharm.github.io/HTA-MELATLAS-1/stories/MEL1-abstract.html)

**Supplementary Minerva story 2.** HTA MEL Atlas 1: Deep Exploration of a Primary Melanoma (https://labsyspharm.github.io/HTA-MELATLAS-1/stories/MEL1-full-story.html)

---

**Figure S1.**
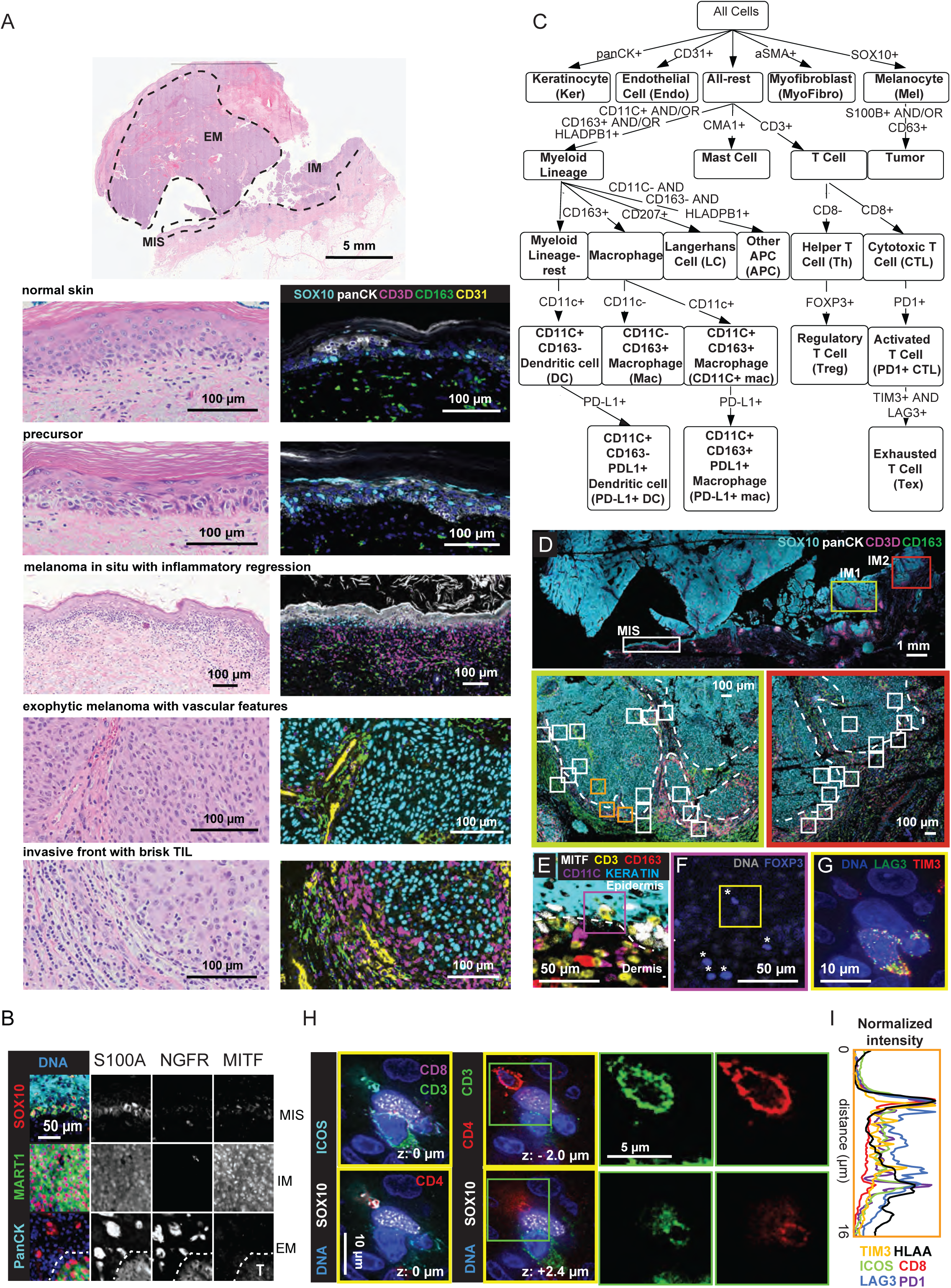
(Related to Figure 1): **(A)** Representative examples of the histopathological features annotated in samples MEL1-MEL13. H&E-stained section of MEL1-1 with three major histologic regions indicated: melanoma in situ (MIS), invasive melanoma (IM), and exophytic melanoma (EM). H&E and the corresponding CyCIF staining of normal, precursor, MIS, IM, and EM regions. **(B)** CyCIF images of MIS, IM, and EM regions (from top to bottom row) of MEL1-1 stained in a composite image (left column) for DNA (blue), the epidermis (PanCK: cyan), and tumor (SOX10: red, MART1: green). Individual grayscale images for staining with S100A, NGFR, and MITF are shown in the right panel. The tumor boundary of EM is indicated by a dotted line. Scale bar, 50 µm. **(C)** Flowchart used in CyCIF experiment 1 indicating strategy used for cell type calling. **(D)** CyCIF of MEL1-1 stained for tumor (SOX10: blue), T-cells (CD3D: purple) and macrophage (CD163: green) markers and epidermis (panCK: white). High-resolution 3D deconvolution microscopy images were obtained from MIS and IM regions. Imaging regions for MIS are presented in panel 1C. Regions within IM1 (green box) and IM2 (red box) are indicated with squares (bottom panel). The invasive front is indicated by a dashed white line. **(E)** CyCIF image of MEL1-1 (same region shown in panel 1G) stained for melanocytes (MITF: white), T cells (CD3: yellow), macrophages (CD163: red), myeloid cells (CD11C: magenta), and keratinocytes (PanCK: cyan). Epidermis, dermis, and dermal-epidermal junction (dashed line) and region of panel 1I (white box) are marked. Scale bar, 50 µm. **(F)** CyCIF of the region shown in panels E and 1G stained for DNA (grey) and Tregs (FOXP3: blue, marked with asterisks). Scale bar, 50 µm. **(G)** Composite high-resolution CyCIF inset from panel F and 1H stained for DNA (blue), SOX10 (white), LAG3 (green), and TIM3 (red). Scale bar, 10 µm. **(H)** Single optical section CyCIF images (same region shown in panel G) depicting the interaction between a Treg (CD4: red and ICOS: cyan), a melanocyte (SOX10: white), and a CTL (CD8: magenta and CD3: green). z=0 µm (left panel; also shown in panel 1I) and z = -2.0 or +2.4 µm (second to left panel; shown also in panel 1J). Magnified regions (green boxes) show co-staining for CD3 (green) or CD4 (red) in each optical section (right panel). Scale bar, 10 µm (left panel) or 5 µm (right panel). **(I)** Quantified spatial distribution of CD8 (red) relative to HLA-A (black) and functional T cell markers ICOS (green), LAG3 (blue), PD1 (magenta), and TIM3 (gold) based on images in panel 1I.

**Figure S2.**
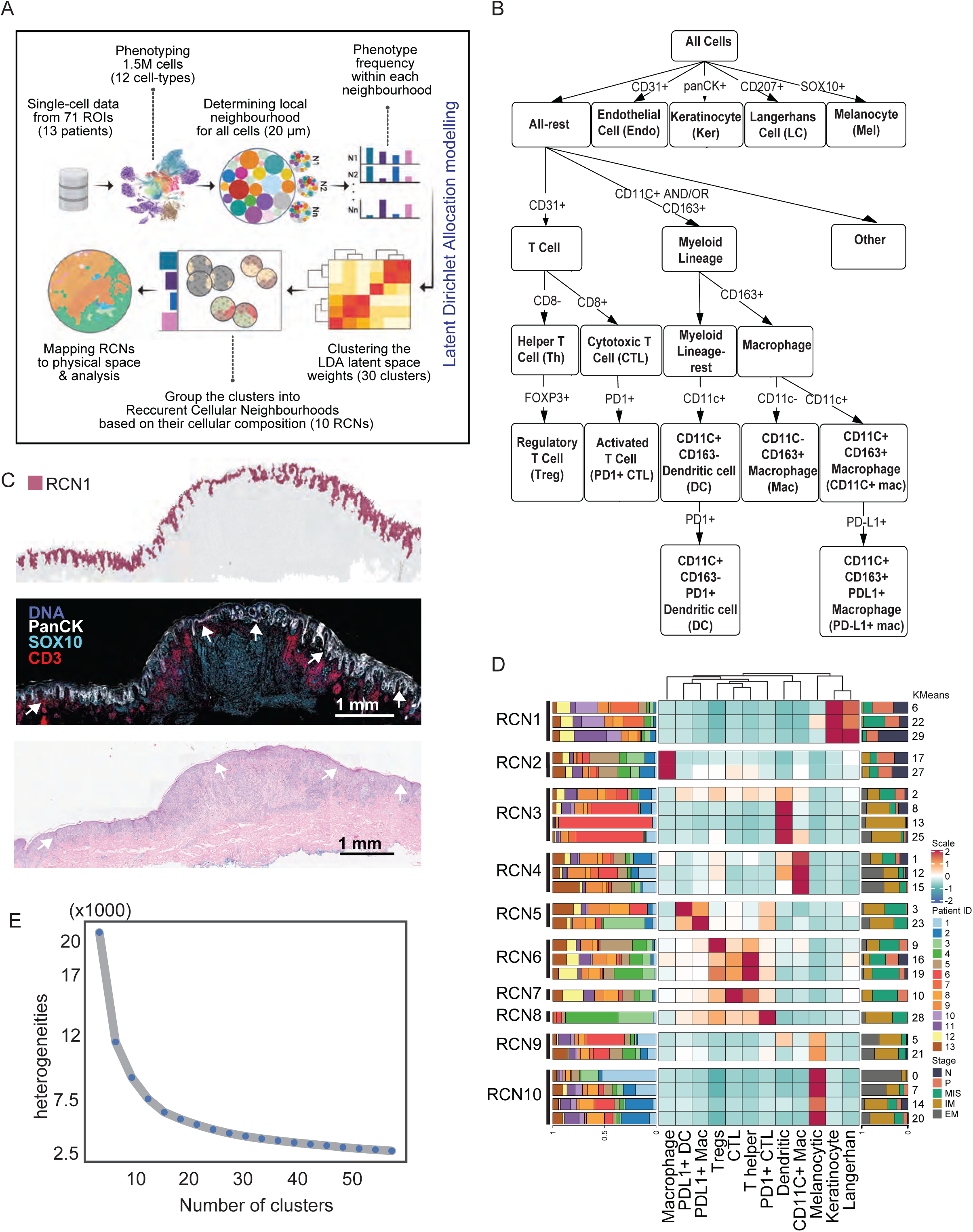
(Related to Figure 2) **(A)** Schematic of the Latent Dirichlet Allocation (LDA) analysis to identify RCNs. Single-cell data from 22-plex CyCIF of 71 ROIs annotated for the stage of melanoma progression from patients MEL1- MEL13 was used for cell type calling to identify 12 distinct cell phenotypes. A spatial-LDA model was trained with a 20 µm proximity radius and the latent weights were subsequently grouped using k-means clustering (k=30) into ten informative meta-clusters (RCN1-10) based on the cellular composition and the frequency of occurrence within the ROIs. **(B)** Flowchart used in CyCIF experiment 2 depicting the strategy used for cell type call. **(C)** Voronoi diagram of RCN1 (top), CyCIF image (middle) with keratinocytes (PanCK: white), melanocytes (SOX10: cyan) and T cells (CD3: red), and the corresponding H&E (bottom) of the same region showing enrichment of RNC1 to the panCK^+^ epidermis (marked with white arrows). **(D)** Heatmap showing the abundance of cell types within the 30 (numbers to the right of the plot) derived cellular neighborhood clusters; as described in methods, a complementary approach was used to that shown in main panel 5D but the results were substantially the same. The clusters are grouped into meta-clusters shown to the left of the plot. The bar chart to the right of the heatmap depicts the distribution of progression stages within each cluster and the left bar chart represents the distribution of patients within each cluster. **(E)** Line plot showing the decay of heterogeneity score with an increasing number of clusters (KMeans) in the x-axis. The latent space vectors of the LDA model were used for generating this plot.

**Figure S3.**
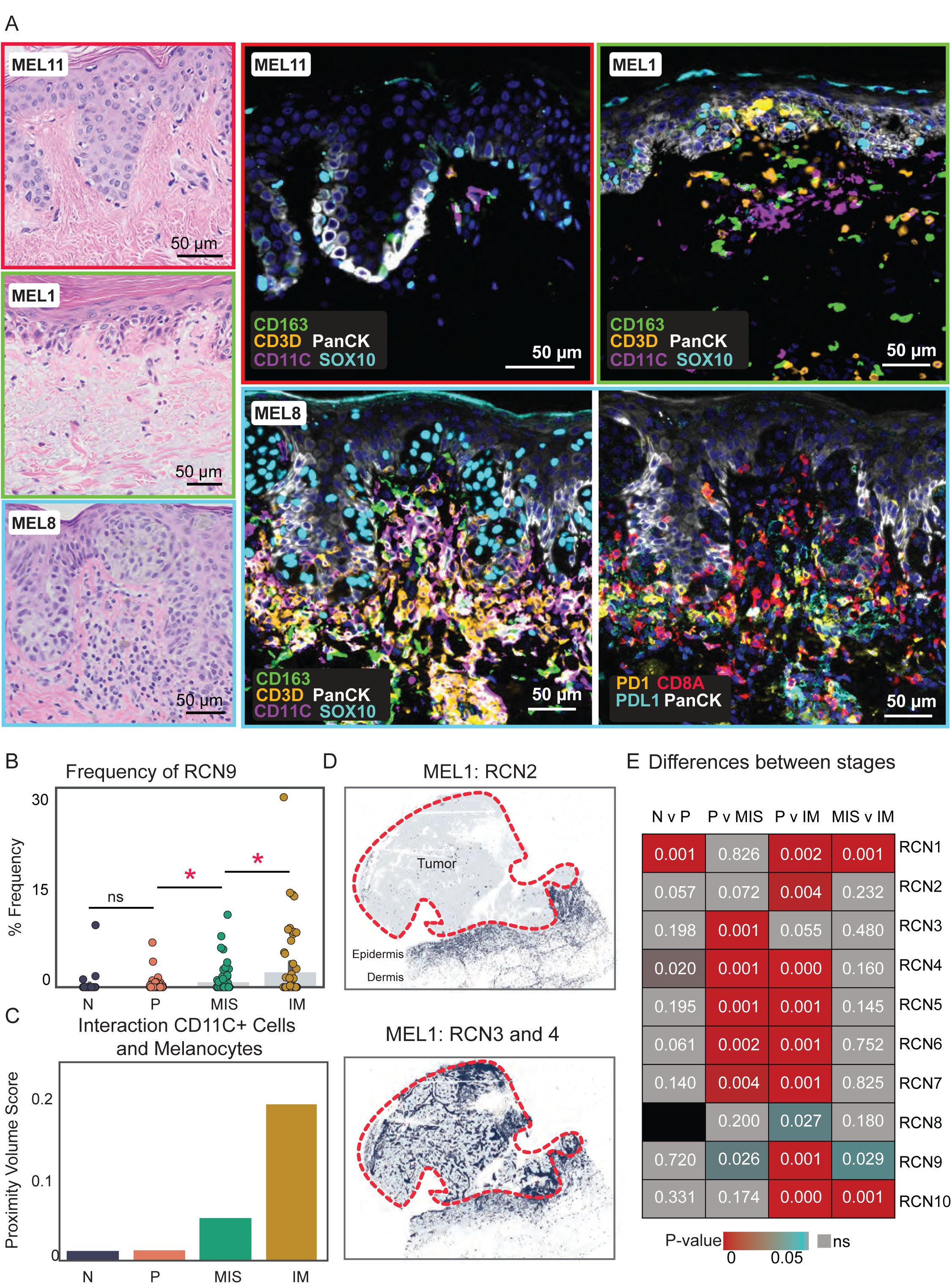
(Related to Figure 3) **(A)** The corresponding H&E images (left column) for normal (MEL11), precursor (MEL1), and MIS (MEL8) CyCIF ROIs that are shown in panel 3C. Four of the CyCIF images presented in panel 3C are magnified in the middle and right columns (MEL11 top left, MEL1 top right, MEL8 bottom panel). The highlighted cell types are melanocytes (SOX10: cyan), keratinocytes (PanCK: white), T cells (CD3: orange), macrophages (CD163: green), dendritic cells (CD11C: purple), PD1^+^ CTLs (PD1: yellow, CD8: red) and PDL1^+^ myeloid cells (PDL1: cyan). Scale bar, 50 µm. **(B)** Swarm plot showing the percent frequency (number of cells belonging to RCN9 divided by the total number of cells within the ROI) of RCN9 between the progression stages. *P <0.05. **(C)** Bar plot of the proximity volume scores across progression stages calculated between CD11C^+^ myeloid cells and melanocytes. **(D)** Scatter plots highlighting cells within RCN2 (top panel) and RNC3 and 4 (bottom panel). RCN2 is spatially restricted to the dermis, while RCN3/4 is more prevalent surrounding the tumor and in the vascular spaces within the tumor. **(E)** Heatmap showing the differences in the frequency of RCN1-10 between progression stages. The significant comparisons (P <0.05) in the RCN frequency between the stages compared are indicated in cyan or red, non-significant comparisons in grey. The comparisons made for each RCN are indicated at the top of the heatmap.

**Figure S4.**
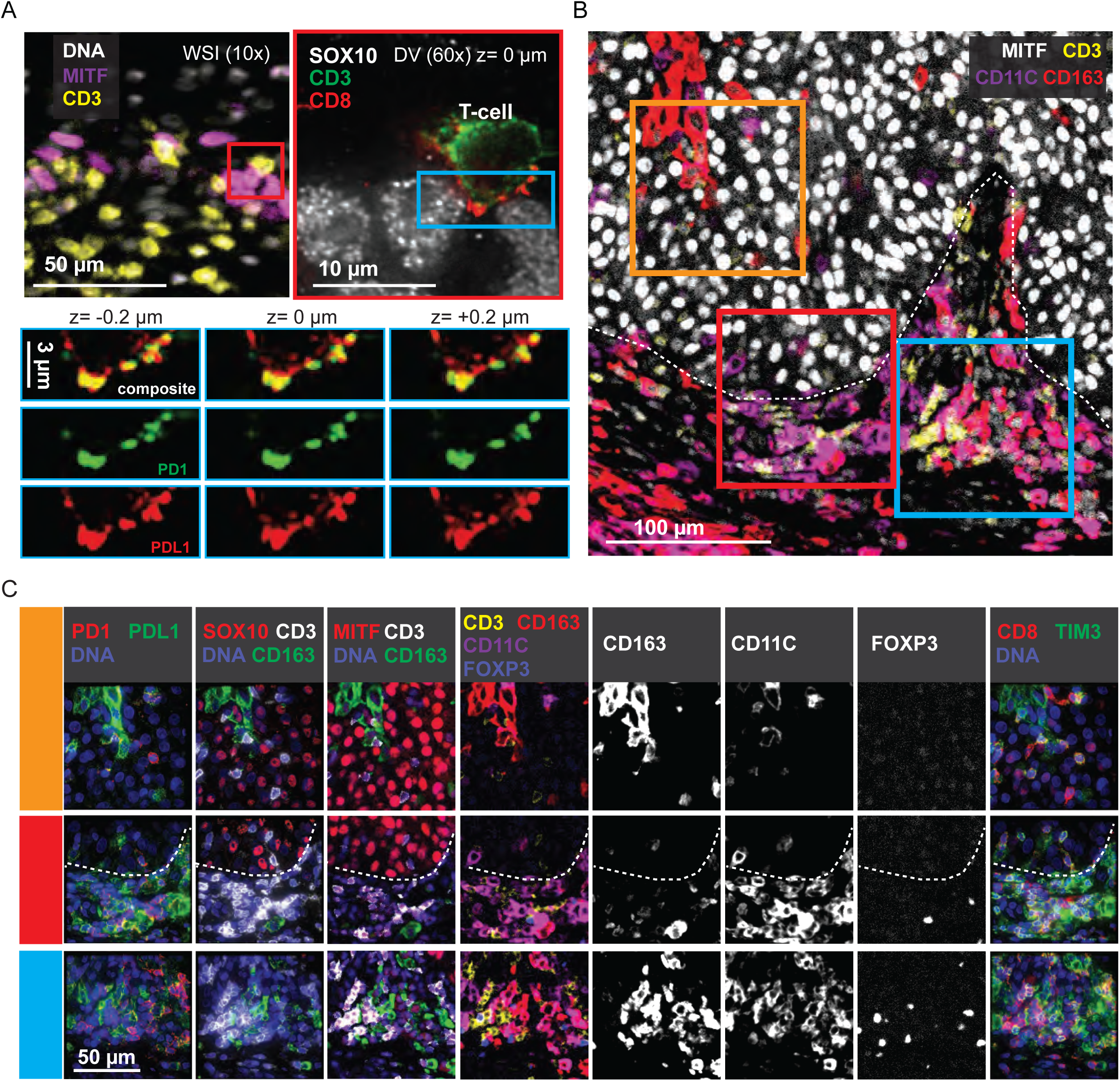
(Related to Figure 4) **(A)** CyCIF image of MIS region of MEL1-1 (top left; same as in panels 1G and S1E), stained for DNA (blue), tumor (MITF: magenta), and T cells (CD3: yellow). A single-optical section (z: 0 µm) of high-resolution CyCIF stained for tumor (SOX10: white) and T cells (CD3: green, CD8: red) is shown in the panel top right (magnification of the inset from the left panel). Below, magnified region of the cell-to-cell interaction (cyan box) with three optical sections (z: -0.2, 0, +0.2 µm) stained for PD1 (red) and PDL1 (green) and a composite image. Scale bars, 50 µm, 10 µm or 3 µm. **(B)** CyCIF image of IM region of MEL1-1 stained for tumor (MITF: white), T cells (CD3: yellow), and myeloid markers (CD163: red, CD11C: magenta). Regions of high-resolution imaging (orange, red, and cyan boxes) and tumor margin (white dashed line) shown in panel C are being indicated. Scale bar, 100 µm. **(C)** CyCIF whole slide (10x; the five panels on the left) and maximum-intensity projection high-resolution (60x; the three panels on the right) images of regions in panel B (orange, red, and cyan boxes). The tumor margin is indicated with a dashed line. Scale bar, 50 µm.

**Figure S5.**
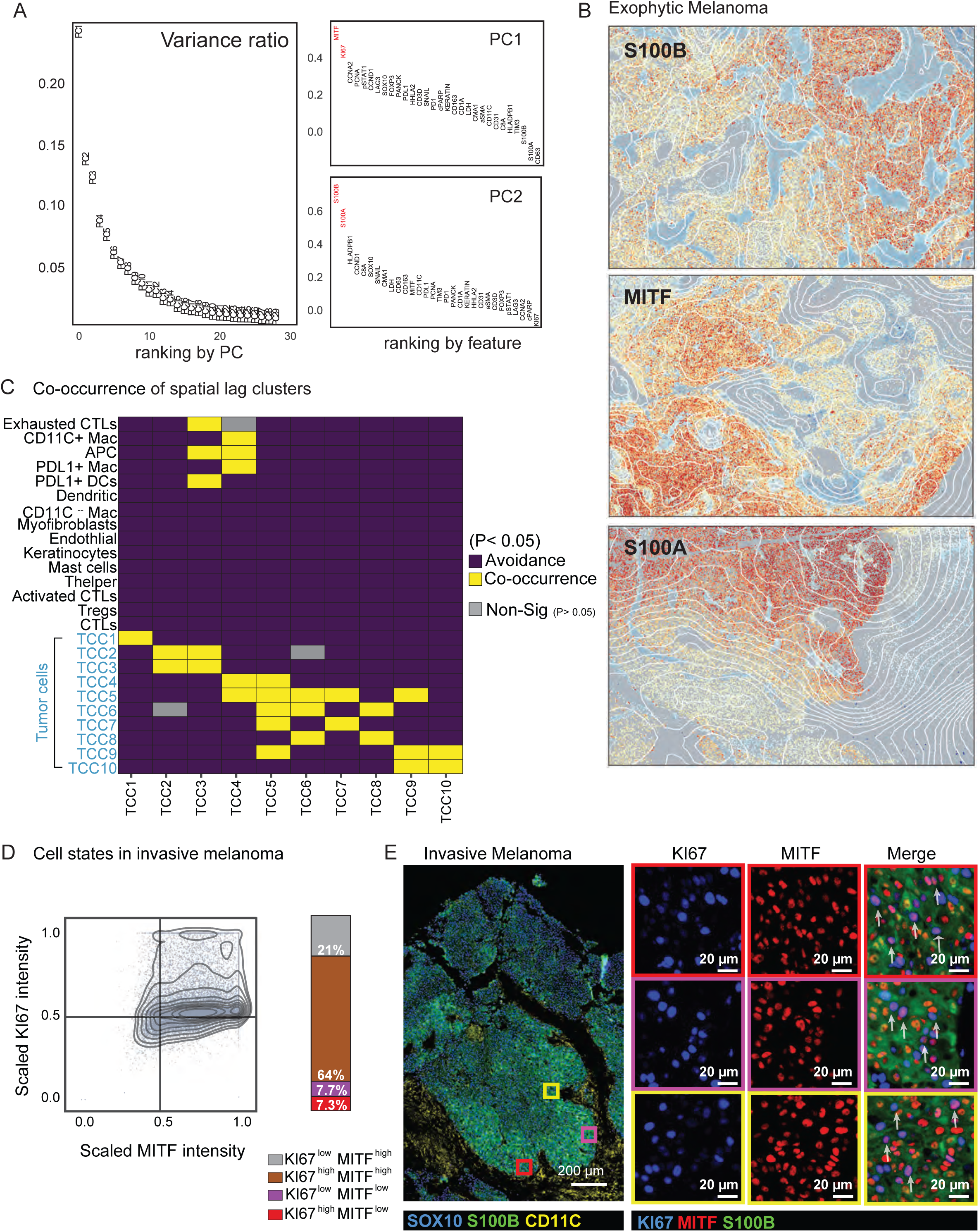
(Related to Figure 5): **(A)** Principal Component analysis variance plot, showing the degree of variance captured by each principal component (PC) within the tumor cells (0.5M cells) in specimen MEL1-1. The plots to the right show the top loadings within PC1 and PC2. **(B)** Regions of EM showing gradient expression patterns of S100B, MITF, and S100A. Contours represent averaged cell expression of the markers and are overlaid on single-cell data. **(C)** Cell-to-cell proximity heatmap showing the presence of significant (P < 0.05) co-occurrence (yellow) or avoidance (violet) between cell types in specimen MEL1-1. **(D)** Scatter plot showing the scaled expression of KI67 and MITF of tumor cells in the IM region. The overlayed contour illustrates the density of the points. The stacked bar graph shows the proportion of cells that falls into each quadrant. **(E)** CyCIF field of view of IM region of MEL1-1 stained for melanocytes (SOX10: blue) and myeloid cells (CD11C: green). The magnified regions indicated with yellow, purple, and red boxes highlight the KI67^+^ (blue) and MITF^+^ (red) tumor cells within these regions. Scale bar, 200 µm (main image) or 20 µm (magnified regions).

**Figure S6.**
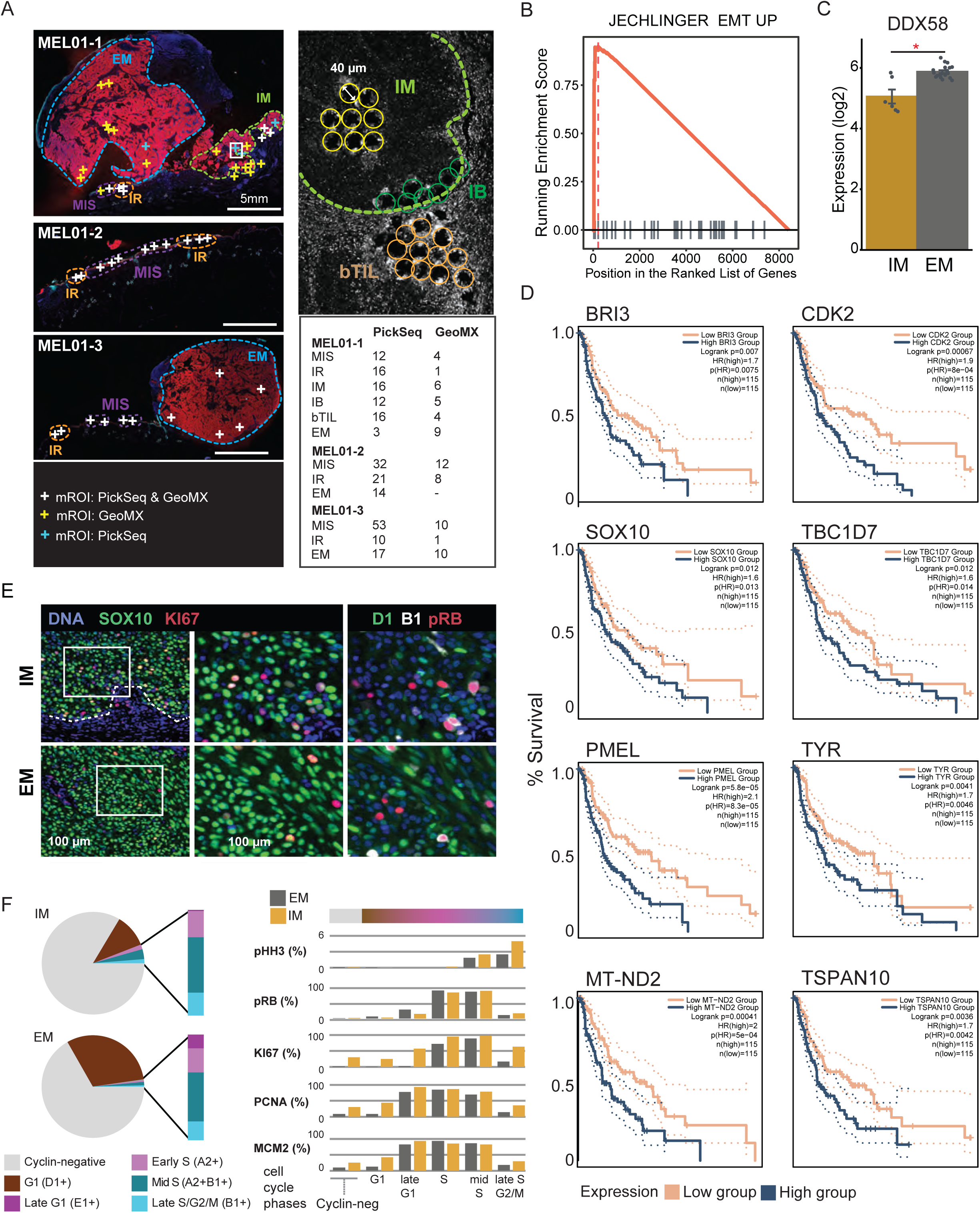
(Related to Figure 6): **(A)** Location of mrSEQ specimens (PickSeq and GeoMX). The left panel shows the whole slide CyCIF images of specimen MEL1-1 to MEL1-3 stained for epidermis (PanCK: cyan) and tumor cells (MART1: red). mrSEQ was performed in areas marked with a color-coded ‘X’ representing regions profiled by GeoMX (yellow), PickSeq (cyan), and both GeoMX and PickSeq (white). The top right panel shows a close-up view of the IM region where mROIs within the tumor (IM), invasive front (IB), and outside the tumor (bTIL) were extracted. The holes show tissue after PickSeq was performed. The bottom right panel shows the number of mROIs extracted by PickSeq and GeoMX between the histological regions. Scale bars, 5 mm. **(B)** Gene set enrichment analysis for upregulation of the EMT pathway in IM (n=16) compared to EM (n=34) mROIs. Derived using PickSeq data. **(C)** Expression of DDX58 between EM and IM mROIs (GeoMX). Data is mean ± SEM; *P <0.05, **P <0.01. **(D)** Kaplan-Meier plots showing survival difference between patients expressing high and low levels of BRI3, CDK2, MT-ND2, PMEL, SOX10, TBC1D7, TSPAN10, TYR in the TCGA melanoma dataset. All highlighted genes showed a significant difference in survival (P <0.05). **(E)** CyCIF image of MEL1-1 IM and EM regions stained for DNA (blue) tumor (SOX10: green), and KI67 (red). (left panel). The magnified regions (white boxes) are also stained for DNA (blue), p21 (green), p27 (red), cyclin D1 (D1: green), cyclin B1 (B1: white), and pRB (red). Scale bars, 100 µm. **(F)** Pie charts (left panel), categorized by exophytic (top) or invasive (bottom) melanoma, showing the percent of tumor cells that stained positive for selected cyclin proteins. Bar plots (right panel) compare the percent of tumor cells in the EM versus IM region that positively stained for phospho-histone H3 (pHH3), phospho-RB1 (pRB), KI67, MCM2, and PCNA (positive: top decile for PCNA or MCM2 staining) as a function of cell cycle: no cyclins (early G1 or G0), cyclin D1 (G1), E2 (G1/S), A2 (early S), A2 co-staining B1 (late S), and B1 alone (G2/M). Co-expression of cyclin D1 with S/G2/M cyclins is depicted in an additional group and suggests a non-cell cycle role for cyclin D1. Cyclin E1-positive cells in the IM region were rare, and this phenotype was not analyzed further.

**Figure S7.**
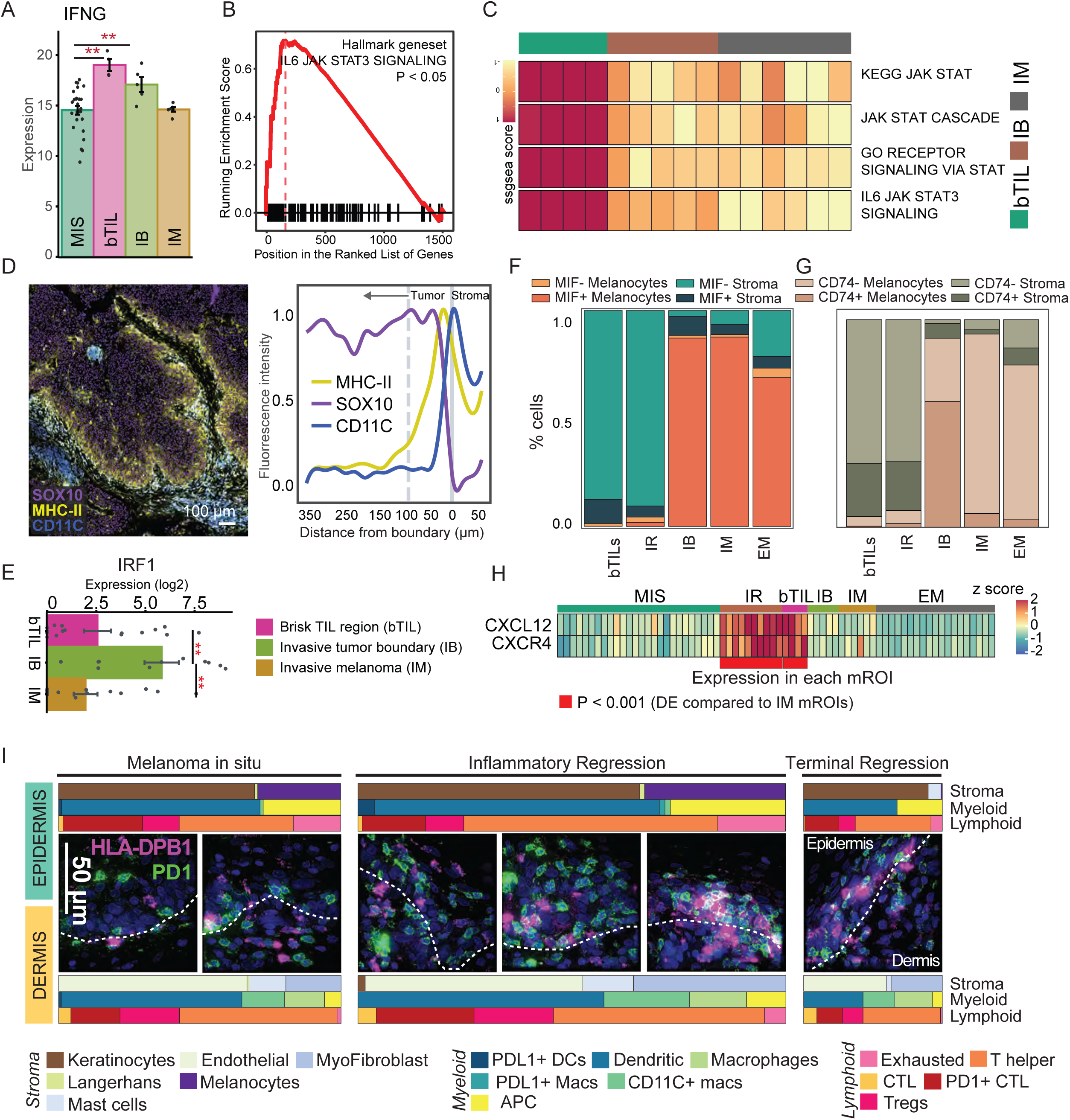
(Related to Figure 7): **(A)** Expression of IFNG in mrSEQ data (GeoMX). Data is mean ± SEM; **P <0.01. **(B)** Gene set enrichment analysis for upregulation of JAK-STAT pathway (from Hallmark gene set) in bTIL region (n=16) compared to tumor regions (EM and IM) (n=50) mROIs. Derived using PickSeq data. P <0.05. **(C)** Single-sample gene set enrichment analysis (ssGSEA) of JAK-STAT related pathways showing enrichment of the pathway in the bTIL region using mrSEQ data (GeoMX). **(D)** CyCIF image of the MEL1-1 invasive tumor front stained for tumor (SOX10: purple), MHC-II (HLADPB1: yellow), and myeloid cells (CD11C: blue) (left). Line plot showing scaled fluorescence intensity of MHC-II (HLADBP1), SOX10, and CD11C within the tumor shown to the left, at, and outside the invasive tumor boundary. The locations of these boundaries are represented by dashed and solid grey lines. Scale bar, 100 µm. **(E)** Expression of IRF1 between bTIL, IB, and IM (PickSeq). Data is mean ± SEM; **P <0.01. **(F)** Stacked barplot showing the proportion of tumor and non-tumor (stromal) cells that are positive or negative for MIF protein expression (CyCIF data) across various histological sites. **(G)** Stacked barplot showing the proportion of tumor and non-tumor (stromal) cells that are positive or negative for CD74 protein expression (CyCIF data) across various histological sites. **(H)** Heatmap showing expression of CXCL12 and CXCR4 (GeoMX). Both genes showed a significant difference in their mean expression (P <0.05) of IR/bTIL compared to IM region. **(I)** CyCIF maximum-intensity projection images of MEL1-1 corresponding to fields of MIS, IR, and terminal regression as indicated in panel 1C. Fields are stained for DNA (blue), PD1 (green), and HLA- DPB1 (magenta). The dermal-epidermal junction, identifying the epidermis and dermis, is indicated with a dashed white line. Scale bar, 50 µm. The bar graph shows the cell type composition for the corresponding histologic regions indicated in panel 1C.

## ACKNOWLEDGEMENTS

We thank David Liu, Genevieve Boland, Jeremy Muhlich, David Weinstock, Robert Krueger, Jared Jessup, and Simon Warchol for their help in multiple stages of this project; we are deeply grateful to Keith Ligon for hosting our clinical research coordinator.

## Notes

https://labsyspharm.github.io/HTA-MELATLAS-1/

